# Engineering, structure, and immunogenicity of a Crimean–Congo hemorrhagic fever virus pre-fusion heterotrimeric glycoprotein complex

**DOI:** 10.1101/2024.04.20.590419

**Authors:** Elizabeth McFadden, Stephanie R. Monticelli, Albert Wang, Ajit R. Ramamohan, Thomas G. Batchelor, Ana I. Kuehne, Russell R. Bakken, Alexandra L. Tse, Kartik Chandran, Andrew S. Herbert, Jason S. McLellan

## Abstract

Crimean–Congo hemorrhagic fever virus (CCHFV) is a tick-borne virus that can cause severe disease in humans with case fatality rates of 10–40%. Although structures of CCHFV glycoproteins GP38 and Gc have provided insights into viral entry and defined epitopes of neutralizing and protective antibodies, the structure of glycoprotein Gn and its interactions with GP38 and Gc have remained elusive. Here, we used structure-guided protein engineering to produce a stabilized GP38-Gn-Gc heterotrimeric glycoprotein complex (GP38-Gn^H-DS^-Gc). A cryo-EM structure of this complex provides the molecular basis for GP38’s association on the viral surface, reveals the structure of Gn, and demonstrates that GP38-Gn restrains the Gc fusion loops in the prefusion conformation, facilitated by an N-linked glycan attached to Gn. Immunization with GP38-Gn^H-DS^-Gc conferred 40% protection against lethal IbAr10200 challenge in mice. These data define the architecture of a GP38-Gn-Gc protomer and provide a template for structure-guided vaccine antigen development.

## INTRODUCTION

Crimean–Congo hemorrhagic fever virus (CCHFV) can cause severe viremia and hemorrhagic fever in humans with case fatality rates up to 40% (1, 2). The virus is spread by ticks of the *Hyalomma* genus, which are distributed widely throughout parts of Europe, Africa, and Asia (3). CCHFV has a broad host range and can infect humans as well as diverse species of wild animals (1). Although transmission to humans mainly results from the bite of infected ticks or contact with infected livestock, human-to-human transmission has also been reported (4, 5). The widening range of the *Hyalomma* tick vector due to climate change, the ability of CCHFV to infect migratory birds and animals subject to livestock trade, the high case fatality rate in humans, and the possibility of human-to-human transmission have resulted in CCHFV being listed as a priority pathogen by the World Health Organization (6). Despite this, no vaccine or therapeutic has yet been approved to treat or prevent infection with CCHFV, highlighting the need for the development of medical countermeasures.

CCHFV belongs to the *Orthonairovirus* genus within the *Nairoviridae* family and *Bunyavirales* order (generally termed nairoviruses and bunyaviruses) (7) and has a negative-sense RNA genome that is made up of small, medium, and large segments. The medium (M) segment encodes for the glycoprotein precursor (GPC), which undergoes several proteolytic processing events in the infected cell to generate immature precursor proteins that are subsequently processed into mature proteins found on the viral envelope (8–10). In addition to a highly glycosylated mucin-like domain (MLD) at the N-terminus, the M segment encodes for three glycoproteins (GP38, Gn, and Gc) and an incompletely understood accessory protein (NSm). The organization of the CCHFV M segment is much more complex than other bunyaviruses, such as hantaviruses, where the M segment encodes for only Gn and Gc (11). As observed for the Gn and Gc proteins of other bunyaviruses, CCHFV Gn and Gc are presumed to form heterodimers that complex into higher-order assemblies on the viral surface (12).

CCHFV Gc is a class II viral fusion protein composed of three domains (I, II, and III) that transition from a metastable pre-fusion conformation to a highly stable post-fusion conformation to facilitate membrane fusion and entry. Within the tip of domain II, Gc contains three fusion loops with hydrophobic residues that insert into the host cell membrane during fusion. Previous structural studies on hantaviruses, which are close relatives of nairoviruses, have demonstrated that the Gc fusion loops exist in distinct pre- and post-fusion conformations (13). For CCHFV, structures have been determined for the post-fusion Gc trimer as well as a pre-fusion-like Gc monomer in complex with two neutralizing antibodies (14, 15). However, in the monomeric CCHFV Gc structure, the fusion loops were in a post-fusion conformation, suggesting that, like hantaviruses, an accompanying glycoprotein stabilizes the Gc fusion loops in the pre-fusion conformation (13, 15).

Gn is the accompanying glycoprotein in bunyaviruses, and CCHFV Gn is generated following SKI-I cleavage from the PreGn precursor at an RRLL cleavage site between GP38 and Gn (8). Gn is implicated in receptor-binding for several bunyaviruses, however, the CCHFV entry factor low-density lipoprotein receptor (LDLR) binds a Gc monomer with high affinity but not monomeric Gn (16–19). Therefore, the function of CCHFV Gn is currently unknown. The structure of CCHFV Gn has not yet been reported, but across the *Bunyavirales* order, structurally characterized Gn proteins exhibit a common domain architecture that CCHFV Gn is expected to share, based in part on a predicted structure generated by AlphaFold2 (20, 21). For example, hantavirus Gn contains three domains within the ‘head’ of Gn that are predominantly composed of β-strands (domain A, β-ribbon domain, and domain B). The ‘base’ is composed of a β-sheet (domain C) followed by an α-helical membrane-proximal external region (13, 20). CCHFV Gn has high structural similarity to hantavirus Gn but is predicted to lack a canonical domain A fold found in most Gn proteins within the *Bunyavirales* order.

Interestingly, GP38, a protein unique to nairoviruses, has been proposed as the domain A equivalent (20). GP38 is released from GP160 and GP85 precursors at an RSKR furin cleavage site between GP38 and the MLD. GP38 has been detected in the supernatant of infected cells, suggesting that the protein is secreted (9). However, there is evidence that suggests GP38 may also localize to the viral surface (22). GP38 in combination with Gn would contain the complete domain architecture observed for Gn proteins of other bunyaviruses, which raises the possibility that GP38 exists at least transiently as a component of a glycoprotein complex with Gn and Gc.

Here, we designed a stable GP38-Gn heterodimer named GP38-Gn^H-DS^, facilitated by an engineered disulfide bond between GP38 and Gn that increases expression and thermostability. A 2.5 Å resolution X-ray crystal structure of the GP38-Gn heterodimer displays a hydrophobically packed GP38-Gn interface. We leveraged the GP38-Gn^H-DS^ design to obtain a 3.4 Å cryo-EM structure of GP38-Gn^H-DS^ in complex with Gc. The complex reveals a GP38-Gn-Gc heterotrimer which is fortified by polar contacts between Gn and Gc, GP38 and Gn, and a contact between GP38 and Gc. The structure of GP38-Gn^H-DS^-Gc provides the basis for GP38’s association with the virion and defines the pre-fusion conformation of the CCHFV Gc fusion loops, which are largely restrained by a highly conserved N-linked glycan on Gn. We also assess the immunogenicity of GP38-Gn^H-DS^-Gc and its ability to protect mice from a lethal CCHFV-IbAr10200 challenge.

## RESULTS

### Prediction and design of a stabilized GP38-Gn construct

To express the GP38-Gn heterodimer, we initially kept the native furin cleavage site (RSKR) between the mucin-like domain (MLD) and GP38 but designed an R516S mutation to abrogate the SKI-I cleavage site between GP38 and Gn, resulting in an uncleaved linker (SRLL, **Supplemental Figure 1A**). This single amino acid substitution boosted expression (**Supplemental Figure 1B**). However, we observed that even when co-expressed with furin protease, SDS-PAGE analysis revealed bands from the purified protein that were consistent with incomplete cleavage of the MLD from the GP38-Gn heterodimer. To address this, we altered the furin cleavage site to six consecutive arginine residues (6xR), which resulted in near complete cleavage of the MLD from the GP38-Gn heterodimer and increased yield of the protein of interest.

We next used AlphaFold2 (21) to predict a model of the full coding sequence of the CCHFV IbAr10200 M segment to facilitate further optimization of the GP38-Gn construct (**Figure 1**). The predicted GP38-Gn heterodimer displayed a conserved domain architecture observed among the Gn and E2 proteins that accompany class II fusion proteins, as described by Guardado-Calvo and Rey (20) and was structurally similar to the Gn protein of Andes virus, the prototypical hantavirus (**Figure 1A**). Based on the domain architecture of CCHFV Gn in the AlphaFold2 prediction and the strategies used to crystallize structures of hantavirus Gn head and base (13), we designed two truncations of the ectodomain construct: one lacking the membrane proximal external region (MPER, residues 520–663, termed GP38-Gn^H+C^) and one lacking both the MPER and Gn domain C (residues 520–587, termed GP38-Gn^H^, **Figure 2A**). Although both truncations increased the expression of the GP38-Gn ectodomain, GP38-Gn^H^ expressed substantially better than either GP38-Gn ectodomain or GP38-Gn^H+C^ (**Figure 2B**).

**Figure 1.**
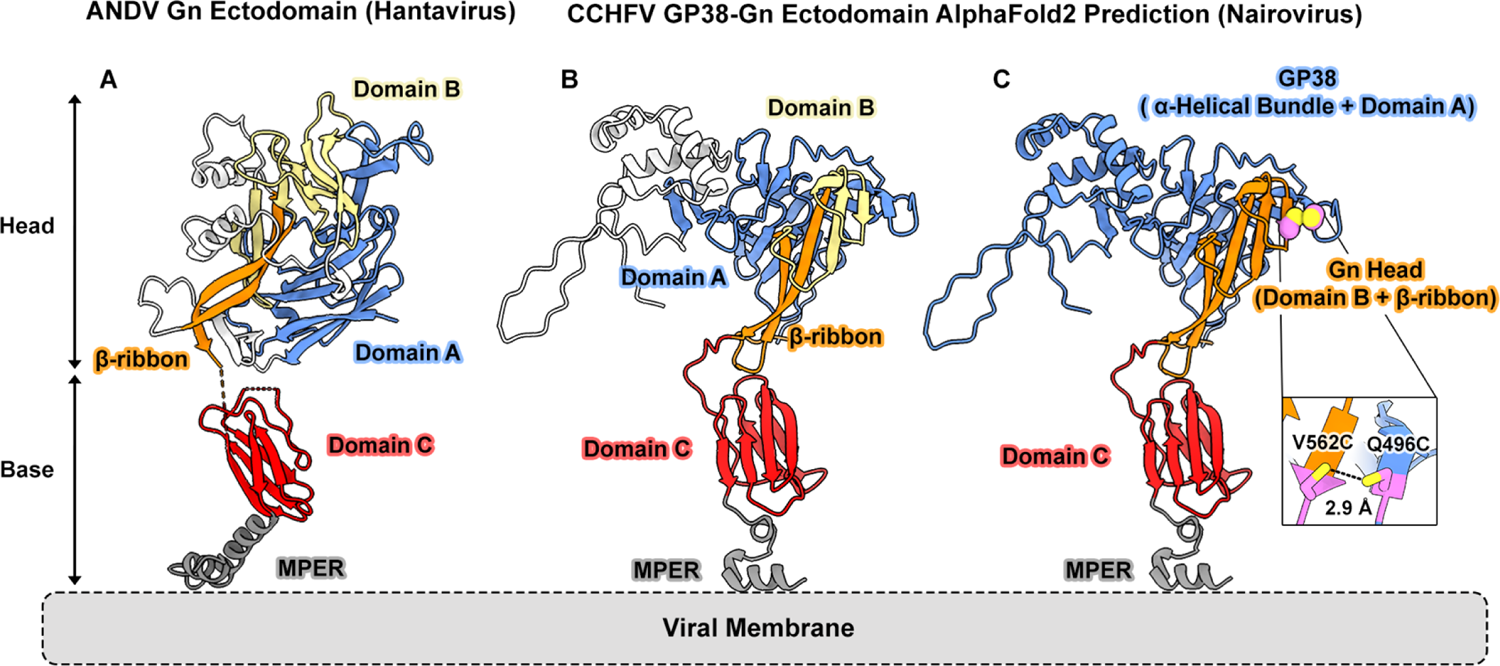
AlphaFold2 predicts a CCHFV GP38-Gn heterodimer with conserved domain architecture characteristic of class II accompanying proteins. A cartoon representation of (A) ANDV Gn (PDB ID: 6ZJM) and (B) an AlphaFold2 prediction of CCHFV GP38-Gn heterodimer (IbAr10200, GenBank AF467768.2) colored by conserved domains of class II accompanying proteins. Domain A is colored in blue, domain B in light yellow, and β-ribbon in orange, and these domains compose the ‘head’ subdomain of the accompanying proteins. Domain C is colored in red and the membrane proximal external region (MPER) in gray. These domains compose the ‘base’ subdomain of the accompanying proteins. Areas without conservation are colored white. (C) AlphaFold2-predicted CCHFV GP38-Gn heterodimer depicted in (B), now colored by GP38 (blue), Gn head (orange), Domain C (red), and MPER (gray) domains. The violet spheres and corresponding inset represent the location of the beneficial cysteine substitutions incorporated into the engineered GP38-Gn heterodimer. Sulfur atoms are displayed in yellow.

**Figure 2.**
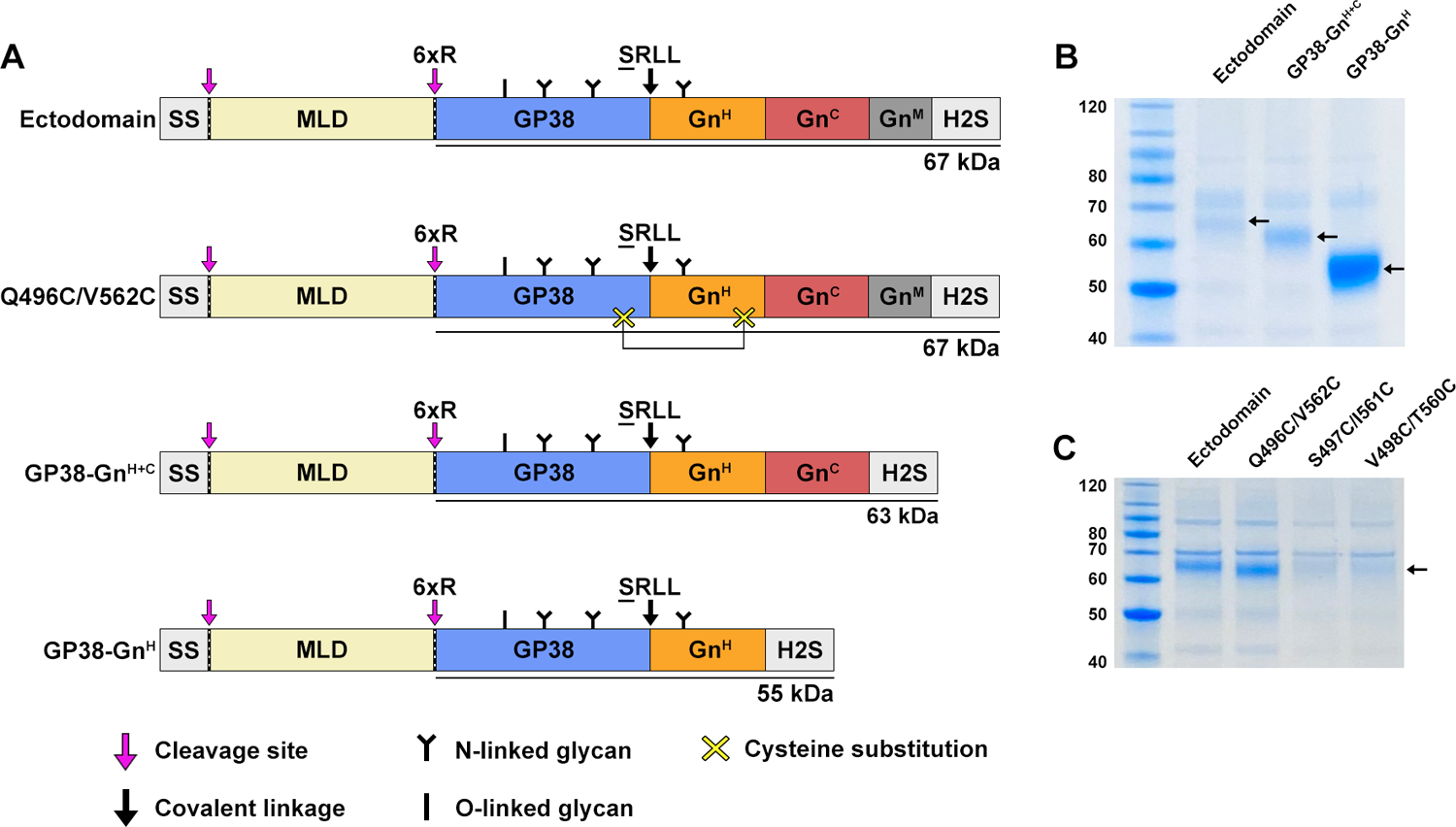
Addition of Q496C/V562C and truncations of Gn boost expression of the GP38-Gn heterodimer. A) A schematic of the GP38-Gn constructs tested for expression. SS represents the native CCHFV M segment signal sequence while H2S represents the HRV3C-cleavable 8x His and Twin-Strep tags. Magenta arrows indicate cleavage sites, and black arrows indicate a mutated cleavage site resulting in a linker with the R◊S mutation underlined. Yellow X’s indicate sites of cysteine substitution, black **Y**s indicate sites of N-linked glycosylation in the mature purified protein of interest, and black **I**s indicate sites of O-linked glycosylation in the mature purified protein of interest. Note that the MLD is highly glycosylated with both N-linked and O-linked glycans, but MLD glycosylation was omitted for simplicity and clarity. (B) SDS-PAGE gel corresponding to relative expression of the GP38-Gn ectodomain construct and two Gn-truncated constructs illustrated in (A). (C) SDS-PAGE gel corresponding to relative expression of the GP38-Gn ectodomain construct and double cysteine mutants in the ectodomain backbone illustrated in (A). Molecular weight standards (kDa) are labeled on the left. Black arrows indicate bands corresponding to the proteins of interest.

To fortify the GP38-Gn heterodimer, we designed three disulfide bonds between the C-terminal β-strand of GP38 and the N-terminal β-strand of Gn domain B and tested these disulfide bonds in the ectodomain construct. Only Q496C/V562C improved expression (**Figure 2C**), with a 2-fold increase. We combined the shortest truncation (GP38-Gn^H^) with the disulfide substitution (Q496C/V562C) to generate our best-expressing construct, GP38-Gn^H-DS^, that expressed at a final yield of 7 mg/L in FreeStyle 293-F cells (**Figure 3A**).

**Figure 3.**
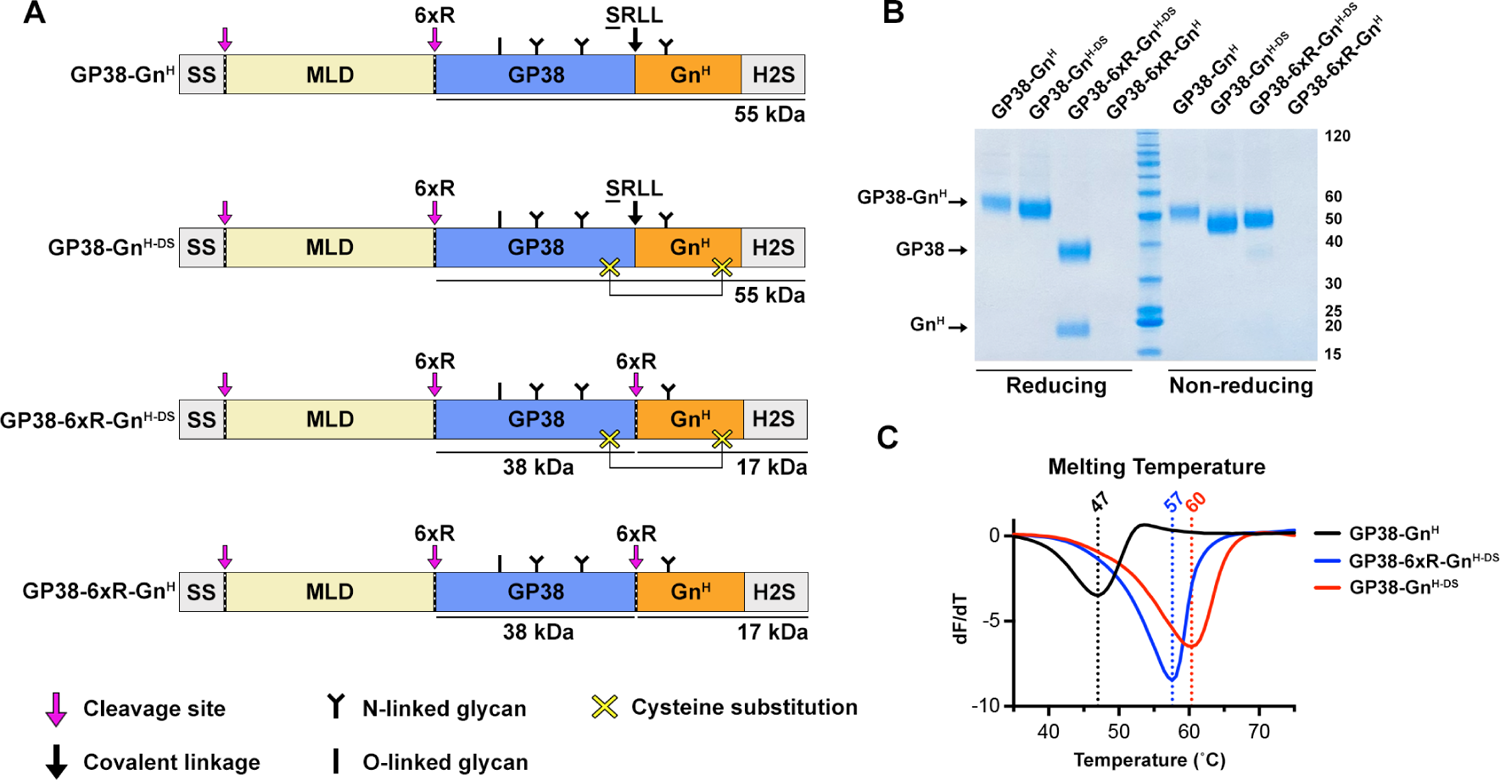
The Q496C/V562C disulfide bond forms and increases T_m_ of GP38-Gn^H^. (A) A schematic of the GP38-Gn^H^ constructs tested for expression. SS represents the native CCHFV M segment signal sequence while H2S represents the HRV3C-cleavable 8x His and Twin-Strep tags. Magenta arrows and dotted lines indicate cleavage sites, and black arrows indicate a mutated cleavage site resulting in a covalent linker with the R◊S mutation underlined. Yellow X’s indicate sites of cysteine substitution, black **Y**’s indicate sites of N-linked glycosylation in the mature purified protein of interest, and black **I**’s indicate sites of O-linked glycosylation in the mature purified protein of interest. Note that the MLD is highly glycosylated with both N-linked and O-linked glycans, but MLD glycosylation was omitted for simplicity and clarity. (B) SDS-PAGE gel corresponding to relative expression the of GP38-Gn^H^ constructs illustrated in (A). Molecular weight standards (kDa) are labeled on the right. (C) Differential scanning fluorimetry thermal stability analysis of expressed variants in (A). The vertical dotted lines and corresponding labels denote the melting temperature for each construct.

To evaluate if the Q496C/V562C mutation in GP38-Gn^H-DS^ formed the intended disulfide bond, we expressed GP38-Gn^H^ and GP38-Gn^H-DS^ variants that replaced the SRLL linker with a 6xR cleavage site and co-expressed the constructs with furin to cleave GP38-Gn at the 6xR junction (**Figure 3A, B**). Introducing the 6xR cleavage site between GP38 and Gn^H^ (GP38-6xR-Gn^H^) eliminated detectable purified protein.

However, the incorporation of Q496C/V562C minimized the deleterious impact of the 6xR cleavage site, as expression of GP38-6xR-Gn^H-DS^ was only slightly decreased relative to GP38-Gn^H-DS^. In agreement with the result in the ectodomain construct, the expression of GP38-Gn^H-DS^ was increased relative to GP38-Gn^H^. As expected, both reducing and non-reducing SDS-PAGE showed a single band for GP38-Gn^H^ and GP38-Gn^H-DS^ at the estimated molecular weight. In contrast, the reducing SDS-PAGE for GP38-6xR-Gn^H-DS^ showed bands at 38 kDa and 17 kDa corresponding to GP38 and Gn^H^, respectively, whereas the non-reducing gel showed a single band at the expected molecular weight for the GP38-Gn^H-DS^ heterodimer. Together, these data confirm that Q496C/V562C results in the successful formation of a disulfide bond between GP38 and Gn.

To evaluate the effect of Q496C/V562C on thermostability, we performed differential scanning fluorimetry (DSF) on GP38-Gn^H^, GP38-Gn^H-DS^, and GP38-6xR-Gn^H-DS^. GP38-Gn^H^ had a melting temperature (Tm) of 47 °C whereas GP38-Gn^H-DS^ had a Tm of 60 °C, indicating that the addition of the disulfide bond increased the Tm by 13°C. Interestingly, replacing the SRLL linker between GP38 and Gn with a 6xR cleavage site decreased Tm by 3 °C, demonstrating that the SRLL linker not only boosted expression, but also increased thermostability of GP38-Gn^H-DS^ (**Figure 3C**). Collectively, these data demonstrate that both the R516S and Q496C/V562C mutations contribute to increased thermostability and enable the production of a GP38-Gn heterodimeric complex.

### A 2.5 Å resolution crystal structure of the GP38-Gn heterodimer reveals a hydrophobically packed interface

Initial attempts to structurally characterize GP38-Gn^H-DS^ via X-ray crystallography resulted in crystals that diffracted poorly. To aid growth of well-diffracting crystals, we created a construct that minimized the flexible loop between GP38 and Gn (residues 509–532) by replacing it with an 8 amino-acid Gly-Ser linker (**Figure 4A**). This construct successfully yielded crystals in space group *P*2_1_2_1_2_1_ that diffracted X-rays to 2.5 Å resolution (**Figure 4B, Supplemental Figure 2**). After iterative rounds of model building and refinement, the final structure had an R_work_/R_free_ of 19.7%/23.7% (**Supplemental Table 1**).

**Figure 4.**
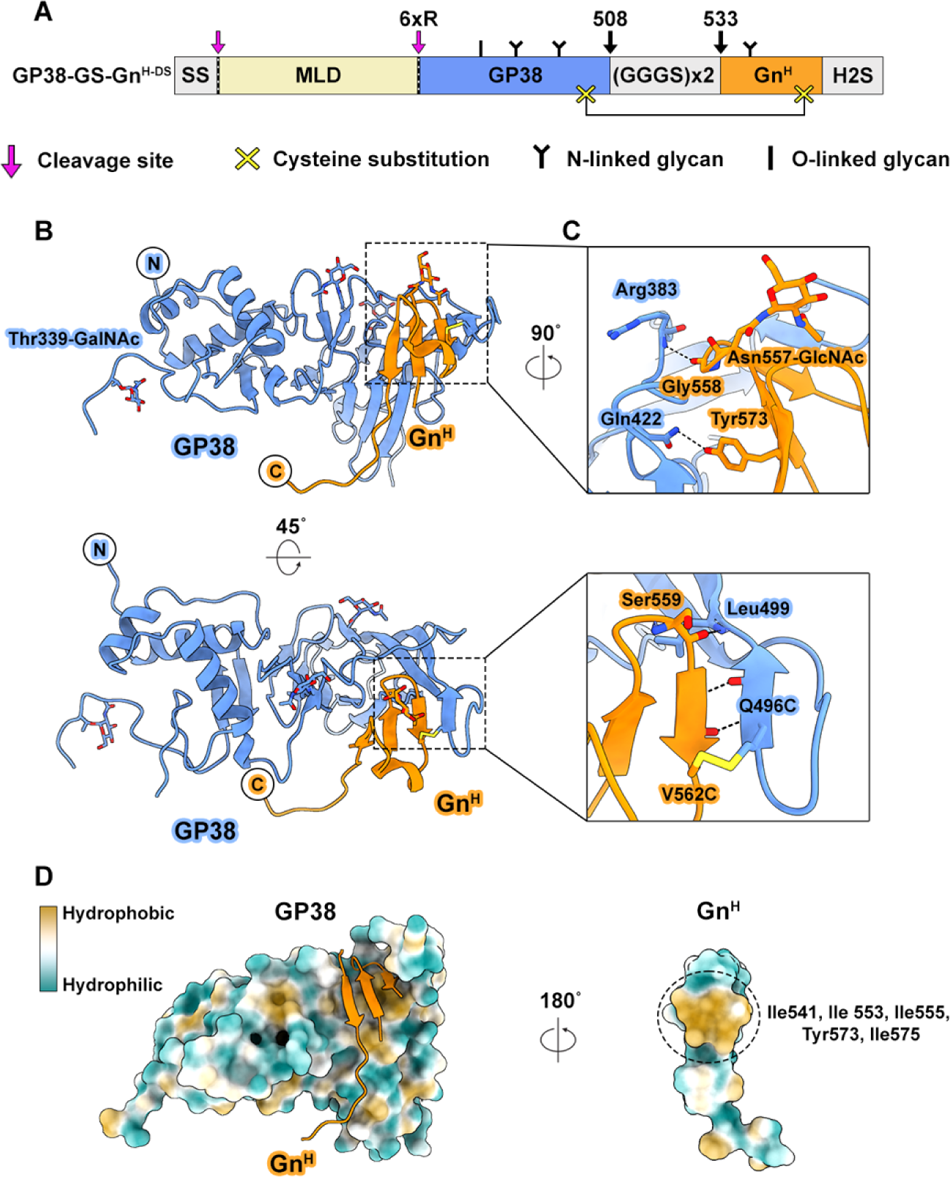
A 2.5 Å resolution crystal structure of GP38-GS-Gn^H-DS^. (A) Schematic of the GP38-GS-Gn^H^ construct. SS represents the native CCHFV M segment signal sequence while H2S represents the HRV3C-cleavable 8x His and Twin-Strep tags. (GGGS)x2 represents the eight amino-acid Gly-Ser linker, and black arrows with the corresponding label indicate the amino acid position preceding and proceeding the Gly-Ser linker (CCHFV IbAr10200 numbering). Magenta arrows and dotted lines indicate cleavage sites. Yellow X’s indicate sites of cysteine substitution, black **Y**’s indicate sites of N-linked glycosylation in the mature purified protein of interest, and black **I**’s indicate sites of O-linked glycosylation in the mature purified protein of interest. Note that the MLD is highly glycosylated with both N-linked and O-linked glycans, but MLD glycosylation was omitted for simplicity and clarity. (B) Side view (upper panel) and top view (lower panel) of GP38-GS-Gn^H-DS^ (PDB ID: 9AVF). GP38 is colored in light blue, and Gn^H^ is colored in orange. N and C indicate the N and C-termini of the protein. (C) Binding interfaces of GP38 and Gn^H^ from the crystal structure. Interacting side chains are depicted as sticks. Hydrogen bonds are depicted as black dotted lines. Oxygen atoms are colored red, nitrogen atoms are dark blue, and sulfur atoms are yellow. (D) A surface view of GP38-Gn^H-DS^ displaying hydrophobicity. The left panel shows GP38 from this crystal structure as a surface view and Gn^H^ from this crystal structure as an orange ribbon. The right panel shows a 180° rotated surface view of Gn^H^. Hydrophobic residues are colored in yellow, and hydrophilic residues are colored in teal.

In the crystal structure, Gn^H^ contains a two-strand domain B fold (residues 550– 572) and an incompletely resolved β-ribbon domain, in which only one of the two strands was observed (residues 573–583 resolved, 533–549 unresolved). Within domain B, we observed an N-linked GlcNAc at Asn557, which is a glycosylation site with 100% conservation among 8 nairovirus sequences analyzed (**Figure 4C, Supplemental Figure 3**). The single GlcNAc observed was expected, as the protein was treated with EndoH to aid in crystallization. In comparison to the previously published crystal structure of the GP38 monomer (23), the conformation of GP38 in the GP38-Gn heterodimer is nearly identical, with a few notable exceptions. In contrast with the monomeric GP38 structure, an O-linked GalNAc on Thr339 is resolved, and we observed a conformational difference in GP38’s C-terminal β-strands (residues 488– 499). These C-terminal β-strands are shifted in the heterodimer to constitute an interface between GP38 and Gn domain B that is further stabilized by the Q496C/V562C disulfide bond. The interface between GP38 and Gn is predominantly hydrophobic and features only a small network of hydrogen bonds, consisting of mostly mainchain-mainchain hydrogen bonds in addition to a contact between the sidechains of GP38 Gln422 and Gn Tyr573. Hydrophobicity analysis highlighted four Ile residues (positions 541, 553, 555, 575) and one Tyr (Tyr573) of Gn^H^ that compose a major hydrophobic packing interface (**Figure 4D**). This hydrophobic patch packs neatly against hydrophobic residues within the interfacing pocket of the GP38 core.

### Immunization with GP38-Gn heterodimers does not elicit neutralizing antibodies

To investigate the immunogenicity of GP38-Gn constructs and their ability to elicit a neutralizing antibody response, we immunized C57BL/6J mice at 0 and 3 weeks with 10 µg of GP38, GP38-Gn^H-DS^ or GP38-Gn^H-DS+C^, adjuvanted with 50 µg poly(I:C).

Mock-immunized mice received PBS, and sera were harvested from all animals at 10 weeks post-immunization (**Figure 5A**). We then quantified the elicited antibody binding titers to GP38, GP38-Gn^H-DS^ and GP38-Gn^H-DS+C^ via ELISA and calculated the area under the curve (AUC) (**Figure 5B**). As expected, the mock-immunized sera bound poorly to the coating antigens (median log_10_AUC values of −1.1 for GP38, −1.6 for GP38-Gn^H-DS^, and −1.2 for GP38-Gn^H-DS+C^). Sera from GP38-immunized mice bound to the GP38 coating antigen with a median log_10_AUC of −0.11 but bound to GP38-Gn^H-DS^ and GP38-Gn^H-DS+C^ less, with values of −0.8 and −0.5, respectively. The sera from GP38-Gn^H-DS^-immunized mice bound to both heterodimeric coating antigens better than the GP38 coating antigen (median log_10_AUC values of −0.8 for GP38, −0.2 for GP38-Gn^H-DS^, and 0.2 for GP38-Gn^H-DS+C^), indicating that immunization with GP38-Gn^H-DS^ elicited Gn-specific antibodies. Interestingly, sera from mice immunized with GP38-Gn^H-DS+C^ bound to all three coating antigens better than sera from mice immunized GP38 or GP38-Gn^H-DS^ (median log_10_AUC values of 0.6 for GP38, 0.8 for GP38-Gn^H-DS^, and 0.8 for GP38-Gn^H-DS+C^). These data demonstrate that GP38-Gn heterodimers are immunogenic and elicit Gn-specific antibodies, and inclusion of Gn domain C increases immunogenicity in comparison to Gn^H^ alone.

**Figure 5.**
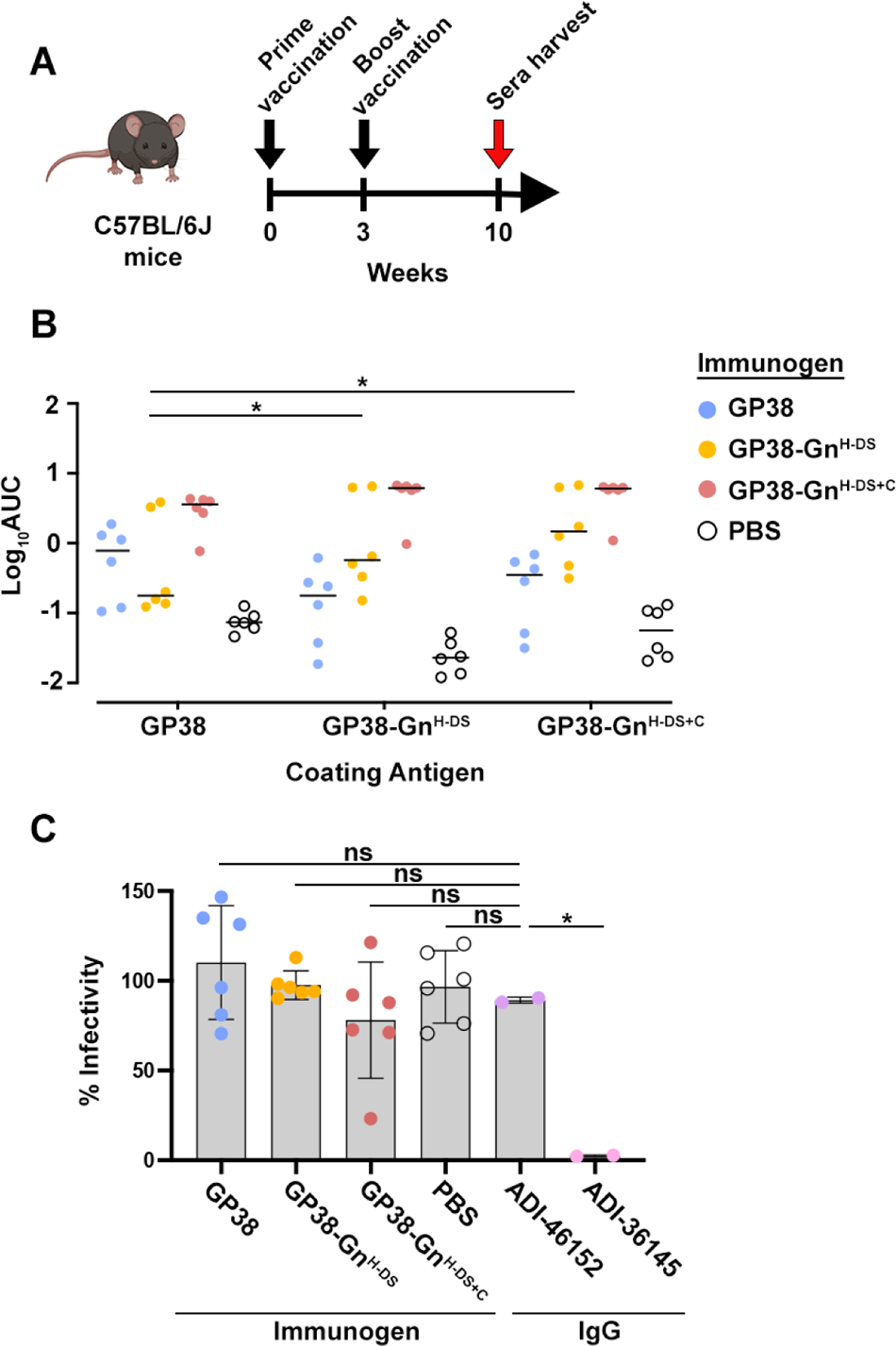
Vaccination of C57BL/6J mice with GP38-Gn constructs. (A) Schematic of the mouse vaccination schedule. Mouse image generated by BioRender. Black arrows indicate vaccination and red arrow indicates sera harvest. (B) ELISA of vaccinated mice sera tested for binding to GP38, GP38-Gn^H-DS^, and GP38-Gn^H-DS+C^. Values shown are the logarithm of the average area under the curve (AUC) for each group. (C) Ability of sera from vaccinated mice to neutralize CCHFV tecVLP infection of Vero cells. Non-neutralizing anti-GP38 antibody ADI-46152 used as a negative control, neutralizing anti-Gc antibody ADI-36145 used as a positive control. Statistical comparison was performed using 2-way ANOVA with Turkey corrections for multiple comparisons (∗∗∗∗p < 0.0001, ∗∗∗p < 0.001, ∗∗p < 0.01, *p <0.05, ns p>0.05).

We next assessed the ability of the sera to neutralize infection of CCHFV transcription- and entry-competent virus-like particles (tecVLPs) (24) (**Figure 5C**). As expected, anti-Gc neutralizing mAb ADI-36145 neutralized infection of Vero cells, whereas anti-GP38 non-neutralizing mAb ADI-46152, the mock-immunized sera, and GP38-immunized sera did not neutralize (25, 26). Additionally, neither GP38-Gn^H-DS^-immunized nor GP38-Gn^H-DS+C^-immunized sera neutralized infection. Therefore, these data highlight a distinct functional difference between the accompanying proteins of hantaviruses (Gn) and CCHFV (GP38-Gn) by demonstrating that GP38-Gn heterodimers do not contain neutralizing epitopes and therefore are not likely involved in receptor binding.

### A 3.4 Å cryo-EM structure of GP38-Gn^H-DS^-Gc reveals a stable heterotrimeric complex in the pre-fusion conformation

To gain a complete structural understanding of the glycoprotein complex, we engineered a construct containing GP38-Gn^H-DS^ in association with the fusion glycoprotein Gc. We hypothesized that, similar to hantaviruses, the Gn ‘head’ subdomain (GP38-Gn^H-DS^ for CCHFV) would make a stable complex with Gc if expressed with a flexible tether between them (13). Therefore, our construct contained the native CCHFV signal sequence and MLD followed by GP38-Gn^H-DS^ joined to Gc (residues 1061–1545) by a Gly-Ser linker. We named this heterotrimeric construct GP38-Gn^H-DS^-Gc (**Figure 6A**). The protein expressed with a yield of 3.5 mg/L in FreeStyle 293-F cells, was monodisperse as judged by size-exclusion chromatography, and had a Tm of 65 °C, which was higher than GP38 (45 °C) or Gc (48 °C) (**Supplemental Figure 4**).

**Figure 6.**
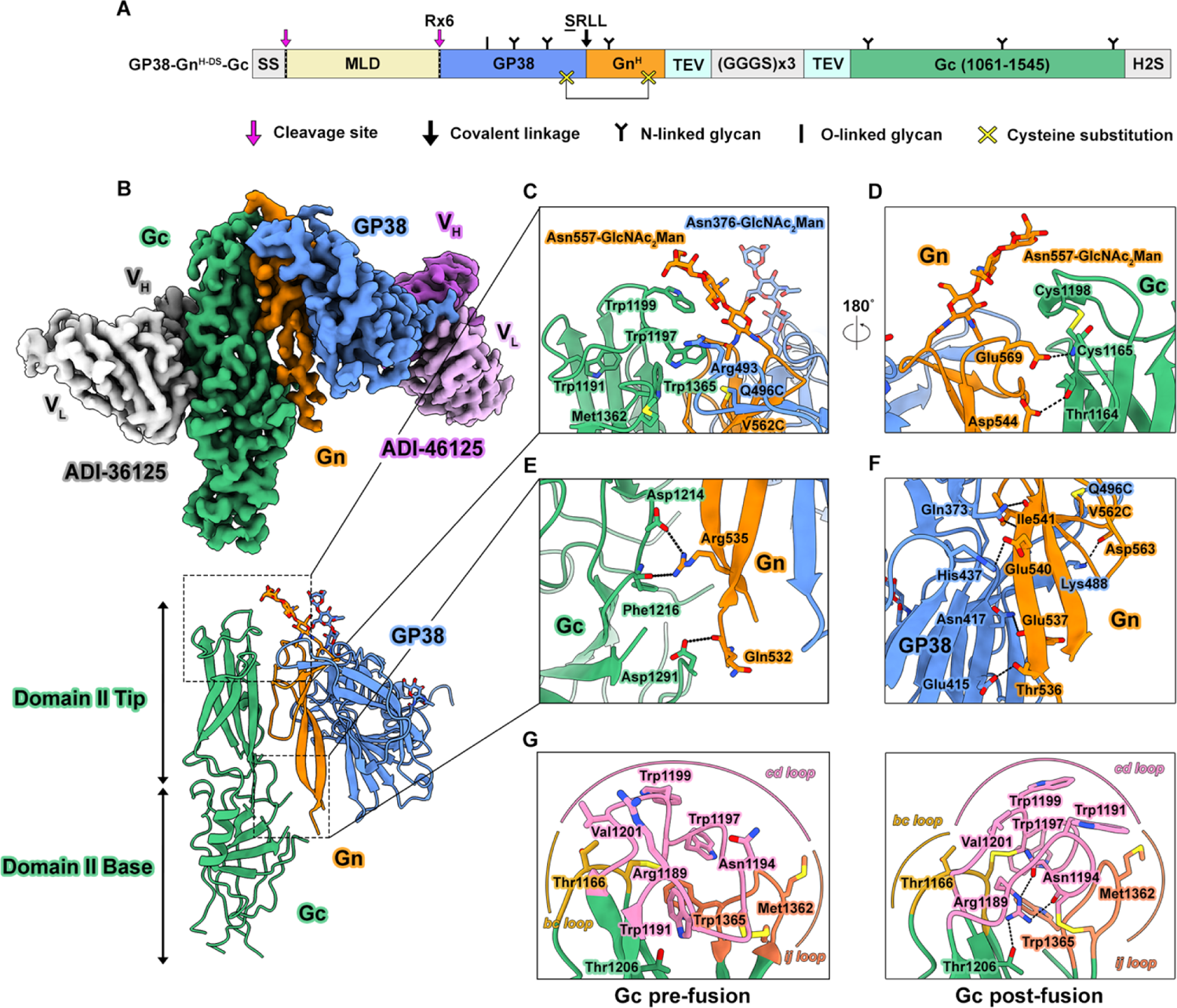
A 3.4 Å resolution cryo-EM structure of GP38-Gn^H-DS^-Gc. (A) Schematic of the GP38-Gn^H-DS^-Gc construct. SS represents the native CCHFV M segment signal sequence while H2S represents the HRV3C-cleavable 8x His and Twin-Strep tags. (GGGS)x3 represents the 12 amino-acid Gly-Ser linker bordered by the TEV cleavage sites in cyan. Magenta arrows and dotted lines indicate cleavage sites, and black arrows indicate a mutated cleavage site resulting in a covalent linker with the R◊S mutation underlined. Yellow X’s indicate sites of cysteine substitution, black **Y**’s indicate sites of N-linked glycosylation in the mature purified protein of interest, and black **I**’s indicate sites of O-linked glycosylation in the mature purified protein of interest. Note that the MLD is highly glycosylated with both N-linked and O-linked glycans, but MLD glycosylation was omitted for simplicity and clarity. (B) Side view of a 3D cryo-EM map of GP38-Gn^H-DS^-Gc bound by ADI-46152 (V_H_ dark purple, V_L_ light purple) and ADI-36125 (V_H_ gray, V_L_ white) Fabs. GP38 is colored in light blue, Gn^H^ is colored in orange, Gc is colored in green. (C) Interfaces of GP38 and Gn^H^ with the fusion loops of Gc and (D) the Domain II tip. (E) Binding interface of Gn^H^ and Gc Domain II base. (F) Binding interface of GP38 and the β-ribbon domain of Gn^H^ (G) Fusion loops of Gc in the pre-fusion (left, PDB ID: 9B8J) and post-fusion (right, PDB ID: 7A59) conformation. Interacting side chains are depicted as sticks. Hydrogen bonds are depicted as black dotted lines. Oxygen atoms are colored red, nitrogen atoms are dark blue, and sulfur atoms are yellow.

Cryo-EM grids of the GP38-Gn^H-DS^-Gc protein in complex with two antibodies were prepared and imaged on a Titan Krios equipped with a K3 detector. Two antibodies—an antigenic site II anti-Gc antibody ADI-36125 and an antigenic site V anti-GP38 antibody ADI-46152—were complexed with GP38-Gn^H-DS^-Gc to increase the mass of the complex and aid particle alignment (25, 26). After curation, a total of 10,739 micrographs were accepted, from which a final stack of 728,031 particles allowed us to obtain a 3.5 Å resolution reconstruction of GP38-Gn^H-DS^-Gc in complex with the two antibodies. To improve resolution in the areas of interest, we performed local refinement using a mask encompassing GP38-Gn^H-DS^, domain II of Gc, and the variable regions of the antibodies, which yielded a 3.4 Å resolution reconstruction (**Figure 6B, Supplemental Figure 5, 6, Supplemental Table 2**).

In the GP38-Gn^H-DS^-Gc model, we observed the same hydrogen bonding interface between GP38 and Gn as seen in the crystal structure of GP38-Gn^H-DS^. However, we observed four additional hydrogen bonds and one salt bridge between GP38 and the β-ribbon domain of GP38-Gn^H-DS^ (**Figure 6F**). GP38 residues Gln373, Glu415, Asn417, and His437 form hydrogen bonds with Gn residues Ile541, Thr536, Glu537, and Glu540 respectively, and GP38 Lys488 forms a salt bridge with Gn Asp563. The newly resolved contacts amend the incomplete characterization of the Gn β-ribbon domain as well as constitute a substantial interface between GP38 and Gn. In total, 1189 Å^2^ of surface area on Gn^H^ is buried by GP38. Surprisingly, we discovered that GP38 not only shares an interface with Gn, but also shares an interface with Gc in the form of a pi-cation stacking interaction between GP38 Arg493 and Gc Trp1197—a key residue in the Gc fusion loops that is indispensable for syncytia formation (**Figure 6C**) (15).

In agreement with the crystal structure of GP38-Gn^H-DS^, the cryo-EM map supported an N-linked glycan attached to Asn557 of Gn. However, in contrast with the single GlcNAc observed in the crystal structure due to the digestion of the branched glycan via EndoH, the cryo-EM map resolved a branched GlcNAc_2_Man (**Figure 6C**).

The GP38-Gn^H-DS^-Gc model contextualizes the significance of this glycan by revealing that the branched glycan shields the *cd* fusion loop of Gc, the epitope of a potently neutralizing antibody ADI-37801 (15). Although CCHFV Gn structurally lacks the capping loop seen in hantaviruses, this glycan, in combination with GP38 Arg493, leaves Trp1197 and Trp1199 relatively inaccessible to solvent. In addition to shielding the Gc fusion loops, Gn makes two polar contacts in the domain II tip: a sidechain-mainchain interaction between Gn Glu569 and Gc Cys1165, and a sidechain-sidechain interaction between Gn Asp544 and Gc Thr1164 (**Figure 6D**). In the domain II base of Gc, the sidechain of Asp1214 and the mainchain of Phe1216 makes hydrogen bonds with the sidechain of Gn Arg535, and the sidechain of Asp1291 makes a hydrogen bond with the mainchain of Gn Thr531 (**Figure 6E**). Together, these data demonstrate that Gn has extensive interfaces with both GP38 and Gc, and GP38-Gn^H-DS^-Gc forms a heterotrimeric complex fortified by many polar interactions.

Hydrophobicity analysis of Gc in the complex revealed two prominent hydrophobic patches: one at the domain II base and one at the domain II tip that is composed of the Trp residues of the fusion loops. While the hydrophobic patch composed of four Ile residues (positions 541, 553, 555, 575) and Tyr573 of Gn was seen in both the crystal structure of GP38-Gn^H-DS^ and the cryo-EM model of GP38-Gn^H-DS^-Gc, we discovered two additional hydrophobic patches on Gn that pack tightly with Gc. In conjunction with the N-linked glycan on Asn557, a hydrophobic patch at the apex of Gn sandwiches the Gc fusion loops, and a hydrophobic patch of residues at the base of Gn^H^ pack against the patch of hydrophobic residues at the base of Gc domain II (**Supplemental Figure 7**).

The Gc analysis also revealed that the fusion loops adopt the pre-fusion conformation, presumably restrained by the presence of GP38-Gn^H-DS^. This is in contrast to the fusion loop conformations that have been previously observed in the structure of monomeric CCHFV Gc in complex with two neutralizing antibodies, as well as structures of the CCHFV Gc post-fusion trimer (14, 15). Comparison of the pre-fusion heterotrimeric GP38-Gn^H-DS^-Gc cryo-EM structure with the published post-fusion homotrimeric Gc crystal structure showed a canonical conformational change in the fusion loops that is comparable to what has been observed in hantavirus Gc structures (**Figure 6G**). Similar to the hantaviruses, the biggest conformational change occurs in the *cd* and *ij* loops, whereas the *bc* loop remains relatively unchanged between the pre- and post-fusion conformations. As expected, from the pre-fusion to post-fusion conformation, Trp1191, Trp1197, and Trp1199 rotate from buried to exposed positions. In contrast to the Trp residues, Asn1194 and Arg1189 transition from exposed to buried positions and form hydrogen bonds with each other. This structural analysis indicates that the presence of Gn, and possibly GP38 as well, is required to restrain the hydrophobic fusion loops of Gc in the pre-fusion conformation.

### Immunization with GP38-Gn^H-DS^-Gc confers 40% protection against lethal IbAr10200 challenge in mice and elicits neutralizing antibodies

Several GP38- and Gc-targeting antibodies are protective in mice, and Gc is the only known target of neutralizing antibodies, highlighting the relevance of both GP38 and Gc in the development of a vaccine antigen (25–27) Because GP38-Gn^H-DS^-Gc contains the glycoproteins relevant in the antibody response against CCHFV and is relatively thermostable in comparison to the monomeric glycoproteins, we hypothesized that immunization with GP38-Gn^H-DS^-Gc would elicit neutralizing antibodies and protect mice from lethal challenge with IbAr10200. C57BL/6J mice were immunized at 0 and 3 weeks with 10 µg of GP38, Gc, GP38 + Gc (5 µg each), or GP38-Gn^H-DS^-Gc, adjuvanted with a 1:1 dilution of Addavax. Mock-immunized mice received PBS. At weeks 0, 3, and 6, sera were harvested for analysis. Mice were challenged with 100 plaque-forming units (PFU) of IbAr10200 on week 7, and one-day post-challenge, mice were transiently immunosuppressed via treatment with 1.5 mg/mouse of IFN-blocking antibody mAb-5A3 (**Figure 7A**). Weight and survival were monitored for 28 days post-challenge. As expected, by the sixth-day post-challenge, all mock-immunized mice succumbed to infection. Interestingly, 40% of mice immunized with GP38, GP38 + Gc, or GP38-Gn^H-DS^-Gc were protected for 28 days after lethal challenge, whereas only 20% of Gc-immunized mice were protected (**Figure 7B**). Notably, mice immunized with GP38 + Gc had improved weight loss recovery and a lower clinical score than all other groups (**Supplemental Figure 8**). These results indicate that, with respect to the surface glycoproteins of CCHFV, GP38 is a key immunogen necessary for the protection against lethal infection, providing 40% protection, and vaccination regimens that include Gc or Gn-Gc did not improve survival.

**Figure 7.**
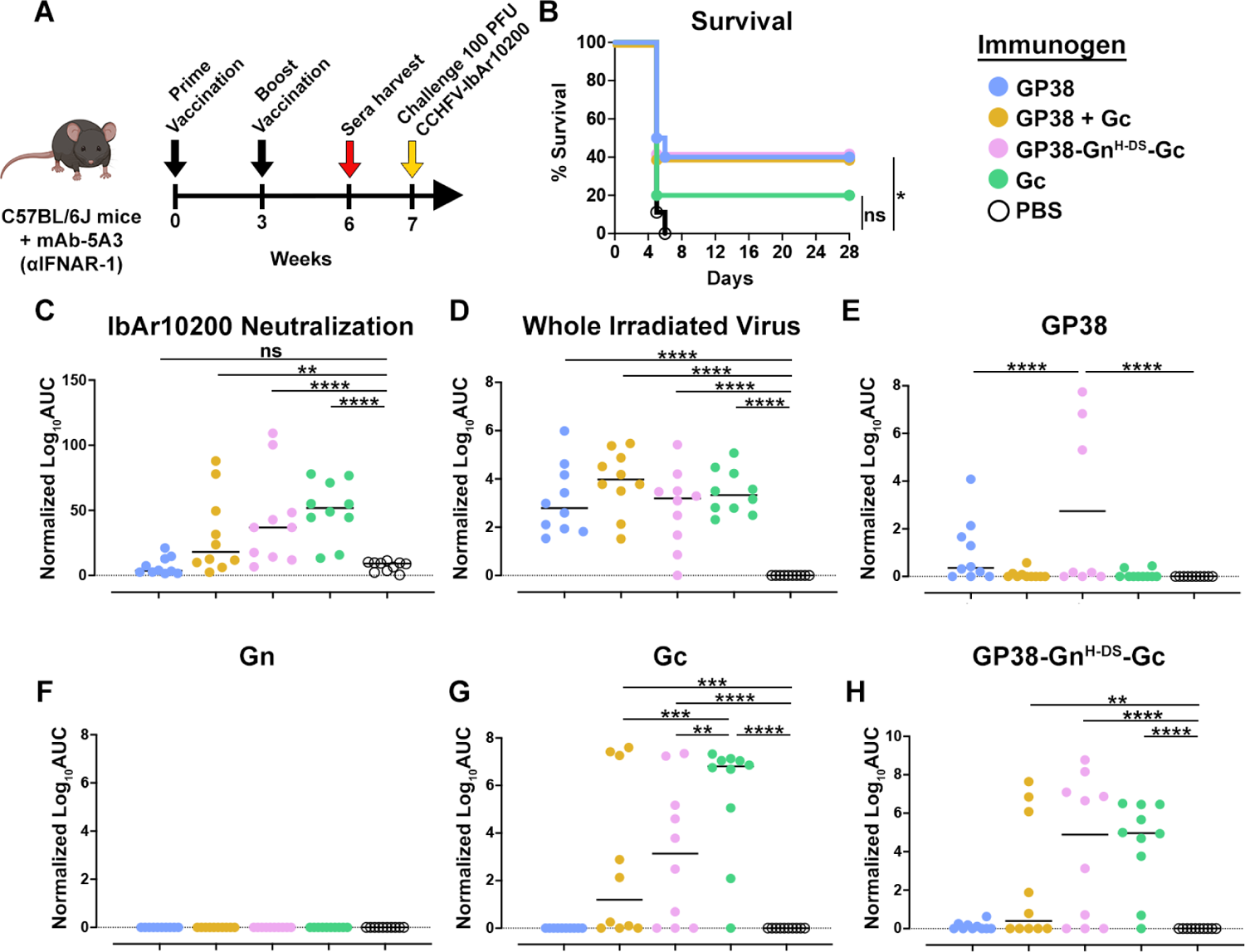
Vaccination of mAb5A3-treated C57BL/6J mice with GPC constructs. (A) Schematic of the mouse vaccination schedule. Mouse image generated by BioRender. Black arrows indicate vaccination, red arrow indicates sera harvest, yellow arrow indicates challenge with IbAr10200. (B) Survival curve of vaccinated mice (n=10). (C) Ability of sera from vaccinated mice (6 weeks post-vaccination) to neutralize IbAr10200 infection of VeroE6 cells. (D-H) ELISA of vaccinated mice sera (6 weeks post-vaccination) tested for binding to whole irradiated virus, GP38, Gn, Gc, and GP38-Gn^H-DS^-Gc. Values shown are the normalized logarithm of the area under the curve (AUC) for each group. Black line reflects the median value. Statistical comparison was performed using 2-way ANOVA with Turkey corrections for multiple comparisons (∗∗∗∗p < 0.0001, ∗∗∗p < 0.001, ∗∗p < 0.01, *p <0.05, ns p>0.05).

We tested the ability of serially diluted sera from vaccinated animals to neutralize IbAr10200 infection of VeroE6 cells (**Figure 7C**) and quantified the AUC. As expected, sera of mock-immunized and GP38-immunized mice did not neutralize infection, whereas sera from Gc-immunized mice neutralized infection, with a log_10_AUC value of 52. GP38 + Gc-immunized mice sera also neutralized infection, with a log_10_AUC value of 18, but did not exhibit a statistically significant difference from the sera of mock-immunized mice. The lower neutralizing capacity is expected as only 5 µg of Gc was used to immunize the GP38 + Gc group. Sera from mice immunized with GP38-Gn^H-DS^-Gc neutralized infection, with a log_10_AUC value of 37, which was statistically different from the mock-immunized control. These results indicate that immunization with GP38-Gn^H-DS^-Gc elicits neutralizing antibodies and the incorporation of GP38, Gn, and Gc in a single chain construct affords an advantage of eliciting higher neutralizing antibody titers than a GP38 + Gc subunit vaccination strategy.

We next evaluated the ability of the sera to bind whole irradiated virus, GP38, Gn, Gc, and GP38-Gn^H-DS^-Gc by measuring binding titers via ELISA and quantifying the AUC. All sera from mice immunized with the CCHFV immunogens bound to whole irradiated virus, and sera from mock-immunized mice did not, confirming that all immunogens elicited CCHFV-specific antibodies (**Figure 7D**). We next evaluated the ability of the sera to bind the monomeric glycoproteins. Interestingly, sera from mice immunized with GP38-Gn^H-DS^-Gc bound to GP38 better than sera from GP38-immunized mice by roughly two logs (**Figure 7E**). None of the sera bound to monomeric Gn, including the sera from mice immunized with GP38-Gn^H-DS^-Gc (**Figure 7F**), indicating no Gn-specific antibodies were detected. Sera from mice immunized with Gc, GP38 + Gc, or GP38-Gn^H-DS^-Gc bound Gc to varying degrees, with sera from Gc-immunized mice showing the strongest binding (**Figure 7G**). We next tested the ability of the sera to bind GP38-Gn^H-DS^-Gc. Surprisingly, sera from GP38-immunized mice did not bind the heterotrimer. Sera from GP38 + Gc-immunized mice bound the heterotrimer but to a lesser extent than the sera from Gc- or GP38-Gn^H-DS^-Gc-immunized mice (log_10_AUC values of 0.4, 5.0, and 4.9, respectively, **Figure 7H**).

Together, these data indicate that immunization with GP38-Gn^H-DS^-Gc elicits neutralizing antibodies, generates a strong GP38-specific antibody response, and produces Gc-specific antibodies with the ability to neutralize infection.

## DISCUSSION

The CCHFV M segment exhibits relatively complex glycoprotein organization and processing, which has hindered the expression of stable protein complexes for structural characterization and vaccine development. In this study, we produced and characterized a stabilized CCHFV glycoprotein complex by first engineering and structurally characterizing a highly thermostable, well-expressing GP38-Gn heterodimer. We then leveraged that design and the structural information to engineer and characterize a prefusion-stabilized heterotrimeric CCHFV glycoprotein complex (GP38-Gn^H-DS^-Gc), which conferred 40% protection against a lethal IbAr10200 challenge *in vivo*.

We found that GP38 forms a stable complex with Gn and Gc (**Figure 6**), therefore establishing that GP38 plays a structural role in a CCHFV glycoprotein complex. Our data support a model whereby GP38 initially folds into a stable component of a heterotrimeric complex before protease cleavage and GP38 dissociation, which is supported by previous studies that detected GP38 in the supernatant of infected cells (9) as well as localized to the viral envelope (22). Both the GP38-Gn^H-DS^ crystal structure and GP38-Gn^H-DS^-Gc cryo-EM structure reported here reveal GP38’s C-terminal β-strands (residues 488–499) in an alternative conformation than in the previously reported structure of GP38 as a monomer (23). This conformational change enables the interaction of GP38 with the Gc fusion loops. The interaction may indicate that GP38 contributes to the pre-fusion stability of Gc, however, the functional roles of GP38 on the viral surface and the extent to which GP38 is present on the mature virion will need to be investigated.

The GP38-Gn^H-DS^-Gc structure defines the architecture of a single GP38-Gn-Gc protomer, but how (or whether) this CCHFV glycoprotein complex organizes into a higher-order assembly remains incompletely understood. The evidence to support a higher-order assembly arises from cryo-ET studies of Hazara virus, a nairovirus closely related to CCHFV, which revealed an ordered and possibly tetrameric assembly (12). Additionally, the structural homology between CCHFV and hantavirus glycoproteins, which assemble into a tetrameric spike lattice, further suggests that nairoviruses may form a tetrameric assembly (13, 20). An understanding of the higher-order assembly of CCHFV glycoproteins may be required to understand the molecular basis of entry and antibody-mediated neutralization, including results obtained with bi-specific antibodies (25), which demonstrated that only certain combinations of Fabs, in a specific organization in the bi-specific, led to enhanced neutralization.

The heterotrimeric structure of GP38-Gn^H-DS^-Gc reveals that GP38 and Gc almost completely mask Gn^H^. Although previous studies have attempted to isolate Gn-specific antibodies from immunized mice or convalescent donors (25, 28), the structural integrity and topological organization of the recombinant proteins used in these experiments have been suggested as rationales for why no Gn-specific antibodies were discovered. Based on the GP38-Gn^H-DS^-Gc structure, probes for antibody isolation that include GP38 with Gn are unlikely to isolate Gn-specific antibodies due to the masking of Gn^H^, rationalizing the results from the previous studies (25, 28). This hypothesis is further supported by the vaccination experiment reported here (**Figure 7**), showing that vaccination with GP38-Gn^H-DS^-Gc did not elicit any detectable Gn-specific antibodies.

Additionally, GP38 may mask Gn^H^ to some extent on the mature virion, which could suggest that if Gn-specific antibodies are raised during natural infection, their abundance is lower in comparison to GP38- or Gc-specific antibodies. In contrast, antibody isolation and characterization experiments for hantaviruses have yielded Gn-specific antibodies that target domain A and domain B (29–32). Antibodies targeting CCHFV GP38 (equivalent to hantavirus Gn domain A) have been isolated (22, 25–27), but a GP38-Gn heterodimer would compose an accompanying protein that is similar in size, organization, and immunogenicity to the Gn protein of hantaviruses.

The results of our immunization experiment further demonstrate that GP38 is a key immunogen, as no survival advantage was observed in mice when including Gn or Gc as antigens (**Figure 7**). These data agree with several studies that have established the importance of GP38 in CCHFV-protection studies performed in murine models (33, 34). Notably, Kortekaas et al. demonstrated that a subunit vaccination with the Gc ectodomain elicits neutralizing antibodies but does not protect mice against lethal challenge (35). However, the dose of Gc administered in those experiments was roughly seven-fold less than what was administered in our study, potentially explaining the increased protection we observed. Although humoral immune responses observed across mice in our vaccination experiment were varied, our construct design and the mutations incorporated into GP38-Gn^H-DS^-Gc may lend itself to improvement of a nucleic acid or viral vectored vaccine by improving the stability and expression of the CCFHV glycoproteins, as well as stabilizing GP38 in the glycoprotein complex.

## ACKNOWLEDGEMENTS

The authors would like to thank Dr. Kaci Erwin for assistance with cell culture and transfection, Christy Hjorth for antibody reagents, Dr. Nicole Johnson for comments on the manuscript, and Drs. Axel F. Brilot and Evan Schwartz at the Sauer Structural Biology Laboratory at the University of Texas at Austin for assistance with cryo-EM data collection. This work was funded in part by the National Institute of Allergy and Infection Disease grant R01AI152246 (J.S.M. and K.C.), the National Institutes of Health grant 1T32GM149364-01 (K.C.), and Welch Foundation grant number F-0003-19620604 (J.S.M.). Use of the NYX beamline 19-ID at the National Synchrotron Light Source II was supported by the New York Structural Biology Center. NYX detector instrumentation was supported by grant S10OD030394 through the Office of the Director, National Institutes of Health. This research used resources of the National Synchrotron Light Source II, a U.S. Department of Energy (DOE) Office of Science User Facility operated for the DOE Office of Science by Brookhaven National Laboratory under Contract No. DE-SC0012704. Additional support from the Structural Biology Center at the Advanced Photon Source.

## AUTHOR CONTRIBUTIONS

Conceptualization: E.M. and J.S.M.; Methodology: E.M. and J.S.M.; Investigation: E.M., S.R.M., A.W., A.R.R., T.G.B., A.I.K., R.R.B., and A.L.T.; Writing – Original Draft: E.M., J.S.M., S.R.M., and A.W.; Writing – Revising and Editing: all authors; Visualization: E.M.; Supervision: J.S.M., K.C., and A.S.H.; Funding Acquisition: J.S.M., K.C., and A.S.H.

## METHODS

### Protein expression and purification of GP38-Gn variants

All variants included the CCHFV M segment (Isolate IbAr10200, GenBank AF467768.2) native signal sequence followed by the MLD, a furin cleavage site (RRRRRR), and GP38-Gn. Codon-optimized (GenScript) variants were cloned into the mammalian expression vector pαH upstream of a human rhinovirus 3C (HRV3C) protease cleavage site, an 8X His tag, and a Twin-Strep tag. GP38-Gn plasmids were transiently co-transfected (4:1 ratio) with a plasmid encoding for a human furin ectodomain (residues 1–794) into 40 mLs of FreeStyle 293-F cells (ThermoFisher). Cells were transfected at a density of 1 million cells/mL using 25 kDa linear polyethylenimine (PEI). Cultures were grown for 6 days in a shaking incubator at 37 °C supplied with 8% CO_2_ and 80% humidity. After 6 days, culture supernatant was separated via centrifugation and passed through a 0.22 μm filter. To evaluate differences in expression, all variants were purified from supernatant using 1 mL of StrepTactin resin following the manufacturer’s instructions (IBA). The elution fractions were concentrated to 100 µL. Purified protein was flash-frozen in liquid nitrogen and stored at −80 °C. Expression experiments of GP38-Gn variants in this study were performed at least twice.

### Protein expression and purification of GP38-Gn^H-DS^-Gc

The GP38-Gn^H-DS^-Gc ectodomain was a single-chain construct that included the native CCHFV M segment signal sequence followed by the MLD, a furin cleavage site (RRRRRR), GP38 (248–515, Q496C), an R516S mutation in the SKI-I cleavage site, resulting in a linker (<u>S</u>RLL, mutation underlined), Gn (520–587, V562C), a Tobacco Etch Virus protease (TEV) cleavable Gly-Ser linker with the sequence **ENLYFQG**GGGSGGGSGGGS**ENLYFQG** (TEV cleavage sites in bold) followed by Gc (1061–1545) upstream of an HRV3C protease cleavage site, an 8X His tag, and a Twin-Strep tag. A codon-optimized (GenScript) gene was cloned into the mammalian expression vector pαH. Expression and purification of GP38-Gn^H-DS^-Gc were performed as described for the GP38-Gn variants. For structural studies, GP38-Gn^H-DS^-Gc was further purified by size-exclusion chromatography (SEC) using a Superdex 200 Increase 10/300 GL column (Cytiva) in a buffer composed of 2 mM Tris pH 8.0, 200 mM NaCl, and 0.02% NaN_3_. Purified protein was concentrated, flash frozen in liquid nitrogen, and stored at −80 °C.

### Protein expression and purification of CCHFV immunogens

The GP38 construct encoded the native signal sequence followed by the MLD, a furin cleavage site (RRRRRR), and GP38. The Gc construct encoded the native signal sequence followed by the MLD, a furin cleavage site (RRRRRR), GP38, another furin cleavage site (RRRRRR), followed by Gc (1061–1545). The GP38-Gn and GP38-Gn^H-DS^-Gc constructs were as described above. Codon-optimized (GenScript) genes were cloned into the mammalian expression vector pαH upstream of a HRV3C protease cleavage site, an 8X His tag, and a Twin-Strep tag. Expression and purification of immunogens were performed as described for the GP38-Gn variants. Immunogens were further purified by SEC using a Superdex 200 Increase 10/300 GL column (Cytiva) in phosphate-buffered saline (PBS). GP38, GP38-Gn^H-DS^, GP38-Gn^H-DS+C^, and GP38-Gn^H-DS^-Gc were used in subsequent ELISA experiments.

### X-ray crystallography

To crystallize GP38-Gn^H-DS^, a construct was expressed that minimized a flexible linker region between GP38 and Gn^H^. This GP38-Gn variant included the signal sequence, MLD, and furin cleavage site as previously described for the other GP38-Gn variants but residues 509–532 (spanning both GP38 and Gn) were replaced with a Gly-Ser linker (GGGSGGGS). To trim the N-linked glycans, the purified protein was brought to a final concentration of 1 mg/mL in a buffer composed of 2 mM Tris pH 8.0, 200 mM NaCl and 0.02% NaN3 and was treated with His-tagged EndoH for 3 hours at a 10:1 ratio of protein to EndoH (wt/wt) at room temperature. During the last hour of EndoH digestion, His-Tagged HRV3C was added at a 50:1 ratio of protein to HRV3C (wt/wt) to remove the affinity tags. Digested protein was passed over Ni-NTA resin to remove cleaved tags, EndoH, and HRV3C. The flow-through was collected, which contained untagged GP38-Gn^H-DS^. The protein was further purified by SEC using a Superdex 200 Increase 10/300 GL column (Cytiva) in a buffer composed of 2 mM Tris pH 8.0, 200 mM NaCl and 0.02% NaN_3_. After concentrating to 7.1 mg/mL, the protein (0.1 µL) was mixed with 0.05 µL mother liquor (8% v/v Tascimate^TM^ pH 8.0, 20% wt/vol polyethylene glycol 3,350) using an NT8 (Formulatrix), and a 0.15 µL drop was spotted onto an MRC2 crystallization tray and sealed to allow for vapor diffusion. The crystal was soaked in mother liquor supplemented with glycerol to a final concentration of 30% (vol/vol), looped, and flash-frozen in liquid nitrogen. Remote data collection was performed at NYX beamline 19-ID at NSLS-II. Diffraction data were indexed and integrated in iMOSFLM (36) before being merged and scaled to 2.5 Å using Aimless (37). Molecular replacement was performed in Phaser (38) using GP38 (PDB ID: 6VKF) as a starting model. The model was built and refined using Coot (39) and Phenix (40). A full description of the data collection and refinement statistics can be found in **Supplemental Table 1**.

### Cryo-EM

0.5 mg/mL GP38-Gn^H-DS^-Gc was incubated with 1.2-fold molar excess of ADI-36125 and ADI-46152 Fab in 2 mM Tris pH 8.0, 200 mM NaCl, 0.02% NaN_3_, and 0.0035% amphipol (wt/vol, Anatrace). The sample was incubated at room temperature for 30 minutes and then deposited onto a glow-discharged Protochips C-Flat 400 mesh 1.2 µm/1.3 µm grid. After 5 sec of wait time, the grid was blotted for 4 sec with a force of 1 using a Vitrobot Mark IV (ThermoFisher) and plunge-frozen into liquid ethane. A total of 13,208 micrographs were collected—8,061 micrographs at 0° tilt and 5,147 micrographs at 30° tilt—from a single grid on a FEI Titan Krios (ThermoFisher) equipped with a K3 direct electron detector (Gatan) at the end of a Gatan Biocontinuum Imaging Filter (Gatan) operated with a 20 eV slit width. The microscope was operated with an accelerating voltage of 300 kV and a total electron flux of 69 e^-^/A^2^. Data were collected at a magnification of 105,000X, corresponding to a calibrated pixel size of 0.83 Å/pix. A full description of the data collection parameters and validation of the structure can be found in **Supplemental Table 2**. CryoSPARC v4.0.1 was used for data processing, which includes patch-based motion correction, patch-based CTF correction, particle-picking, and then curation of the particles via iterative rounds of 2D classification. 3D volumes were generated using *ab initio* reconstruction, and data were further processed via heterogeneous refinement, homogenous refinement, and subsequently non-uniform homogeneous refinement of final classes. The processing workflow can be found in **Supplemental Figure 5**. Models were docked into the experimental EM map using ChimeraX (41) and further refined by Phenix and Coot. The starting models for Gc and ADI-46152 were from PDB ID 7L7R and 8VWW. For GP38 and Gn, the starting model was the crystal structure from this manuscript (PDB ID: 9AVF). For ADI-36125, an initial homology model was generated using the SAbPred server (42).

### Differential scanning fluorimetry

All variants were prepared at a concentration of 3 µM with a final concentration of 10X SYPRO Orange Protein Gel Stain (Sigma-Aldrich) in a white, opaque 96-well plate. 15 fluorescence measurements (λex = 465 nm, λem = 580 nm) were taken per °C using a Roche LightCycler 480 II, with a temperature ramp rate of 0.04 °C/sec, and a temperature range of 22 °C to 95 °C. Data were plotted as the derivative of the melting curve. Each curve is an average of three technical replicates.

### Cell culture

Vero and VeroE6 cells, immortalized epithelial cell lines isolated from the kidney of an adult female African grivet monkey (RRID:CVCL-0059 and RRID: CVCL-0574, respectively), were obtained from the American Type Culture Collection (ATCC). SW13 cells, a cell line isolated from the adrenal gland and cortex of a 55-year old female patient with carcinoma (RRIDD:CCL-105), were obtained from ATCC. BSR-T7 cells (RRID: CVCL_RW96), generated by stable T7 RNA polymerase expression in BHK-21 cells, were a kind gift from K.-K. Conzelmann. The parent cell line (RRID: CVCL_1915) was isolated from the kidney of a 1-day-old male golden hamster. All cells were cultured in Dulbecco’s modified Eagle’s medium (DMEM, high glucose; Thermo Fisher Scientific) supplemented with 10% heat-inactivated fetal bovine serum (FBS; Gibco), 1% penicillin-streptomycin (P/S; Thermo Fisher Scientific), and 1% GlutaMAX (Thermo Fisher Scientific). All cell lines were maintained in a humidified 37°C incubator supplied with 5% CO_2_.

### *In vivo* immunization with GP38-Gn constructs

For immunization with GP38-Gn constructs, 6–8-week-old female C57BL/6J mice were purchased from the Jackson Laboratory (JAX: 000664). Mice (n =6) were immunized with 10 µg of GP38, GP38-Gn^H-DS^, GP38-Gn^H-DS+C^, or an equal volume of phosphate-buffered saline (PBS) adjuvanted with 50 µg of poly(I:C) (High Molecular Weight) VacciGrade (InvivoGen) and boosted three weeks after in the same fashion. Constructs and adjuvant were each diluted in endotoxin-free PBS (Millipore) to a total volume of 100 µL each before they were mixed to obtain a total of 200 µL per mouse. 200 µL of immunogen + adjuvant was delivered by intraperitoneal (IP) injection for each mouse.

Mice were bled two days before prime (day −2) and ten weeks post-boost (day 70). Sera were isolated from whole blood by allowing the blood to coagulate for 1 hour at room temperature and separated by microcentrifugation (10000 *g*, 10 min). Sera were aliquoted and stored at −80 °C for subsequent use in enzyme-linked immunosorbent assay (ELISA) and tecVLP neutralization assays.

### ELISAs for GP38-Gn immunization

ELISAs for each serum sample were run in duplicate. Flat bottom, high binding, half-area 96-well plates (Corning) were coated with 75 ng of each immunogen in PBS at 4°C overnight. The initial coating was decanted, and the plate was then blocked with 5% nonfat dry milk (BioRad) in PBS for 3 hours at 37 °C. Serial 3-fold dilutions were made for each mouse serum sample starting at a 1:100 dilution in 3% bovine serum albumin (Fisher) in PBS and incubated with the coated immunogens for 1.5 hours at 37 °C. Sera/immunogen samples were incubated with an anti-mouse IgG-HRP secondary (Jackson Immunolabs; Cat. 115-035-003) in 3% nonfat dry milk in PBS for 1 hour at 37°C. Finally, the ELISA was developed for 8 min at room temperature with 1-Step TMB Ultra Substrate (Thermo) before being neutralized with 0.5 M H_2_SO_4_. Binding was quantified using Cytation 5 cell imaging multimode reader (BioTek) to measure absorbance at 450 nm. After background subtraction, area under the curve (AUC) values were computed for each ELISA curve (Baseline Y =0, Ignore peaks <10% of the distance from min to max Y). AUCs were log-transformed to fulfill normality and homoscedasticity requirements, and log-transformed AUCs for each serum and immunogen group were compared by 2-way ANOVA with Tukey corrections for multiple comparisons. Statistical tests were performed in GraphPad Prism 10.0.3.

### Generation of tecVLPs bearing CCHFV IbAr10200 glycoproteins

The amino acid sequence for the IbAr10200 was derived from GenBank M segment sequences with an accession number NC_005300. Transcription- and entry-competent virus-like particles (tecVLPs) bearing CCHFV glycoproteins were generated as previously described (24, 25). Briefly, BSR-T7 cells were transfected with plasmids encoding the CCHFV nucleoprotein (NP), glycoprotein complex (GPC), polymerase (L), as well as T7 polymerase, and a minigenome encoding Nano-Glo Luciferase. 15 hours post-transfection, transfection media was replaced on cells with fresh DMEM growth media. 48 hours post-transfection, tecVLP-containing supernatants were collected, clarified by low-speed centrifugation, and finally pelleted by ultracentrifugation at 25,000 rpm for 2.5 hours. Pelleted tecVLPs were resuspended in plain DMEM overnight before storage at −80 °C prior to use in any infection experiments.

### tecVLP neutralization assay

Vero cells were seeded in 96-well cell culture plates (Corning) at 18,000 cells per well 24 hours before infection. The number of tecVLPs to add was empirically determined such that the maximum luminescence signal of infected cells was ∼500x that of background. Mouse sera were incubated with tecVLPs at either a 1:20 or 1:100 dilution in DMEM supplemented with 2% FBS, 1% P/S, and 1% GlutaMAX for 1 hour at 4 °C. 90 µL of media was removed from the target cells, and 50 µL virus/sera mixture was in duplicate. The infection was allowed to proceed 14-16 hours at 37 °C and 5% CO_2_. The infection media was then decanted, cells were washed once with PBS, and the luciferase signal was developed using the Nano-Glo luciferase assay system (Promega) per the manufacturer’s instructions. Infectivity was quantified by luminescence signal using Cytation 5 cell imaging multimode reader (Biotek) with the following run parameters: 200 gain, 6 mm height, 10 sec integration time. Percent infectivity values were calculated for each serum sample relative to sera-free, infected wells. Sera group infectivity values were assessed for normality and were compared by 2-way ANOVA against the control group (mock-immunized) with Turkey corrections for multiple comparisons (GraphPad Prism 10.0.3).

### Ethics statement for IbAr10200 *in vivo* challenge study

Murine challenge studies were conducted under Institutional Animal Care and Use Committee (IACUC)-approved protocols in compliance with the Animal Welfare Act, PHS Policy, and other applicable federal statutes and regulations. The facility where these studies were conducted (USAMRIID) are accredited by the Association for Assessment and Accreditation of Laboratory Animal Care, International (AAALAC) and adhere to the principles stated in the Guide for the Care and Use of Laboratory Animals, National Research Council, 2013. Humane endpoints were utilized during these studies and mice that were moribund, according to an endpoint score sheet and in line with IACUC-approved criteria, were humanely euthanized.

### IbAr10200 *in vivo* challenge study

4-week-old male and female C57BL/6J mice (strain #000664; The Jackson Laboratory) were vaccinated two times at 3 week intervals with 10 µg of recombinant protein or PBS control diluted at a 1:1 ratio with Addavax (InvivoGen) via the IP route. Constructs were diluted in endotoxin-free PBS (ThermoFisher Scientific) to a total volume of 200 µL each before they were mixed with 200 µL Addavax to obtain a total of 400 µL per mouse.

Whole blood was collected prior to vaccination on days 0, 21, and 42 by submandibular bleed and sera were isolated from whole blood as described above. Mice were challenged on day 53. For the challenge, all mice were challenged with 100 plaque-forming units (PFU) of CCHFV-IbAr10200 by the IP route. 24 hours post-challenge, mice were transiently immunosuppressed by treatment with 1.5 mg/mouse of mAb-5A3 (Leinco Technologies Inc.) via the IP route. Mice were monitored daily for weight changes, clinical score, and survival. Mice were scored on a 4-point grading scale; 1 defined by decreased grooming and ruffled fur, 2 defined by subdued behavior when un-stimulated, 3 defined by lethargy, hunched posture, and subdued behavior even when stimulated, and 4 defined by bleeding, unresponsiveness, severe weakness, or inability to walk. All mice scoring a 4 were considered moribund and were euthanized based on IACUC-approved criteria. Daily observations were increased to a minimum of twice daily while mice were exhibiting a clinical score of 3.

### Viruses

The authentic CCHFV isolate CCHFV-IbAr10200 was used in this study.

### Neutralization assays against authentic CCHFV

Neutralization assays were conducted as described previously (25). In brief, heat-inactivated serum was diluted 1:5, and then serial 3-fold dilutions were generated. CCHFV-IbAr10200 was incubated with dilutions for 1 hour at 37 °C. The serum-virus mixture was added to monolayers of VeroE6 cells in a 96-well plate at a final multiplicity of infection (MOI) of 0.06 and incubated for 1 hour at 37 °C. Infection medium was then removed, and fresh cell culture medium without sample was added. 48 hours post-infection, culture medium was removed, and plates were fully submerged in 10% buffered formalin and fixed for at least 4 hours at room temperature. Plates were removed from formalin and permeabilized with 0.2% Triton-X for 10 minutes at room temperature and treated with blocking buffer (Cell Stain Buffer; ThermoFisher). Infected cells were detected by consecutive incubation with CCHFV-specific antibody 9D5 (3 µg/ml; BEI NR-40270) and secondary detection antibody (goat anti-mouse) conjugated to AlexaFluor 488 (1:2000 dilution; Invitrogen). Percent infection was determined using the Cytation5 high-content imaging instrument and data analysis was performed using the Gen5.11 software (BioTek). Percent infectivity values were calculated for each serum sample relative to sera-free, infected wells and normalized to naïve control samples. AUC values were computed for each neutralization curve (Baseline Y=0, Ignore peaks <10% of the distance from min to max Y) and log transformed. AUCs were compared by 2-way ANOVA with Turkey corrections for multiple comparisons (GraphPad Prism 10.0.3).

### Preparation of irradiated CCHFV whole antigen

SW13 cells were infected with CCHFV-IbAr10200 at an MOI of 0.01 and incubated for 1 hour at 37 °C. After incubation, infection medium was removed, and fresh cell culture medium was added. 72 hours post-infection, supernatants from CCHFV-IbAr10200 infected SW13 cells were harvested and cleared from cell debris through load-speed centrifugation at 2,000 rpm for 10 minutes at 4 °C. Clarified supernatants were centrifuged at 12,000 rpm for 4 hours at 4 °C to pellet virus and pelleted virus resuspended in cold 1x Tris-NaCl-EDTA (TNE) buffer (Quality Biologics). Virus was loaded onto a 20-60% sucrose gradient and spun at 40,000 rpm (SW-41 rotor) for 16 hours at 4 °C with no brake. Virus band was collected and treated by gamma-irradiation (8×10^6^ rads).

### Enzyme-linked immunosorbent assay (ELISA)

High bind half-area ELISA plates (Corning) were coated overnight (∼18 hours) at 4 °C with either 125 ng CCHFV rGc, Sheep Fc-Tag (Native Antigen), 125 ng rGn, His-Tag (Native Antigen), 75 ng GP38 protein, 75 ng GP38-Gn^H-DS^-Gc, or 125 ng irradiated CCHFV whole antigen diluted in PBS per well. The following day, plates were blocked with 5% skim milk (BD Biosciences) diluted in PBS containing 0.05% Tween-20 (PBST) for 2 hours at 37 °C. Sera samples were diluted 1:10 with subsequent 3-fold dilutions (dilution range 1:10 to 1:7290) in blocking buffer and plates were loaded with dilutions in duplicate. Plates were incubated at ambient temperature for 2 hours, washed three times with PBST, and then incubated with horseradish peroxidase (HRP) conjugated goat anti-mouse (1:2000 dilution; Jackson ImmunoResearch) diluted in blocking buffer for 1 hour at ambient temperature. After incubation, plates were washed three times with PBST and then developed with TMB substrate (ThermoFisher Scientific). Reaction was stopped using 0.16 M sulfuric acid and absorbance was read at 450 nm wavelength, detected using a SpectraMax (Molecular Biosciences) microplate reader.

Naïve sera collected prior to vaccination was used as an internal control for each group. A cutoff value was determined based on the average absorbance of the naïve control starting dilution plus 3 standard deviations. Only sample dilutions whose average were above this cut-off were registered as a positive signal. AUC values were computed for each ELISA curve (Baseline Y=0, Ignore peaks <10% of the distance from min to max Y). AUCs were log-transformed and compared for each serum and immunogen group by 2-way ANOVA with Tukey corrections for multiple comparisons. Statistical tests were performed in GraphPad Prism 10.0.3.

### Quantification and statistical analysis

Statistical details, including the number of replicates (n), measures of precision, and the statistical test used for each experiment can be found in the corresponding figure legends and in the results section. All area under the curve values (ELISA and microneutralization assays) were log transformed prior to statistical analysis. All statistical analyses were conducted in GraphPad Prism 10.0.3.

---

**Supplemental Figure 1.**
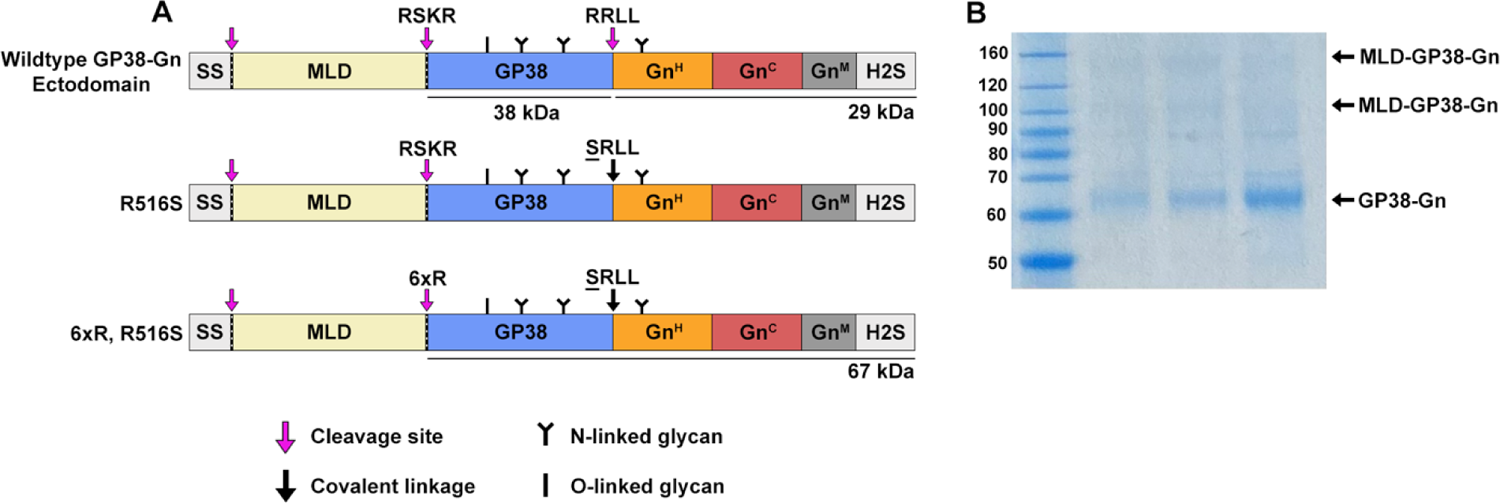
R516S boosts expression of GP38-Gn ectodomain and six consecutive arginine residues increase yield of protein of interest. (A) A schematic of the GP38-Gn constructs in panel B. SS represents the native CCHFV M segment signal sequence while H2S represents the HRV3C-cleavable 8x His and Twin-Strep tags. Magenta arrows and dotted lines indicate cleavage sites, and black arrows indicate a mutated cleavage site resulting in a covalent linker with the R→S mutation underlined. Black **Y**’s indicate sites of N-linked glycosylation in the mature purified protein of interest, and black **I**’s indicate sites of O-linked glycosylation in the mature purified protein of interest. Note that the MLD is highly glycosylated with both N-linked and O-linked glycans, but MLD glycosylation was omitted for simplicity and clarity. (B) An SDS-PAGE of GP38-Gn constructs. Molecular weight standards in kDa are on the left.

**Supplemental Figure 2.**
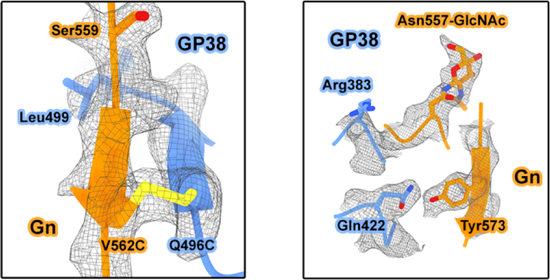
Electron density of stabilizing disulfide bond, Asn557-GlcNAc, and interfacing residues in the GP38-Gn^H-DS^ crystal structure. Electron density (2Fo-Fc) is shown as gray mesh. GP38 is shown in blue and Gn is shown in orange. Side chains are depicted as sticks. Oxygen atoms are shown in red, nitrogen atoms are shown in dark blue, and sulfur atoms are shown in yellow.

**Supplemental Figure 3.**
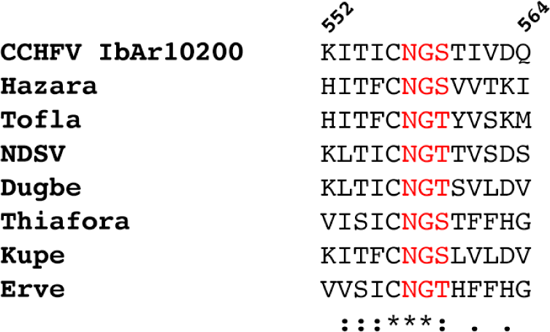
The Gn Asn557 glycosylation site is highly conserved across nairoviruses. Sequence alignment of residues 552–564 (IbAr10200 numbering) of nairovirus Gn. Multiple sequence alignment was generated by Clustal Omega. Asterisks indicate positions that have fully conserved residues, colons indicate strong conservation among groups, and periods indicate weak conservation among groups. Red text indicates N-X-T/S N-linked glycosylation site.

**Supplemental Figure 4.**
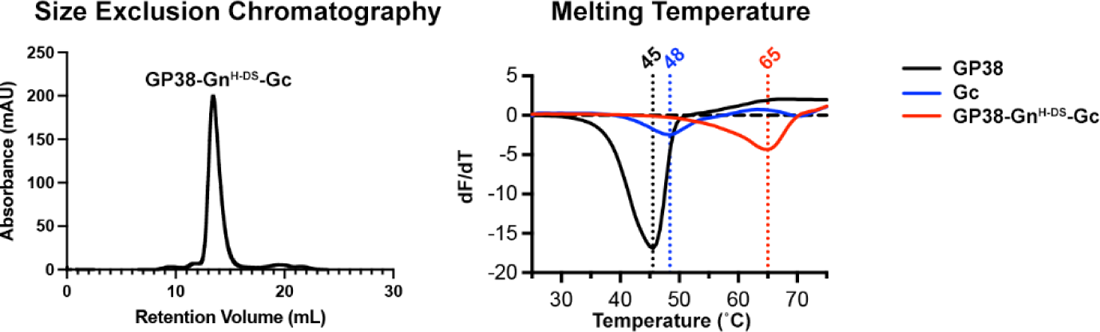
SEC and DSF of GP38-Gn^H-DS^-Gc. Size exclusion chromatography for GP38-Gn^H-DS^-Gc (left). Differential scanning fluorimetry thermal stability analysis of GP38, Gc, and GP38-Gn^H-DS^-Gc (right). The vertical dotted lines and corresponding labels denote the melting temperature for each construct.

**Supplemental Figure 5.**
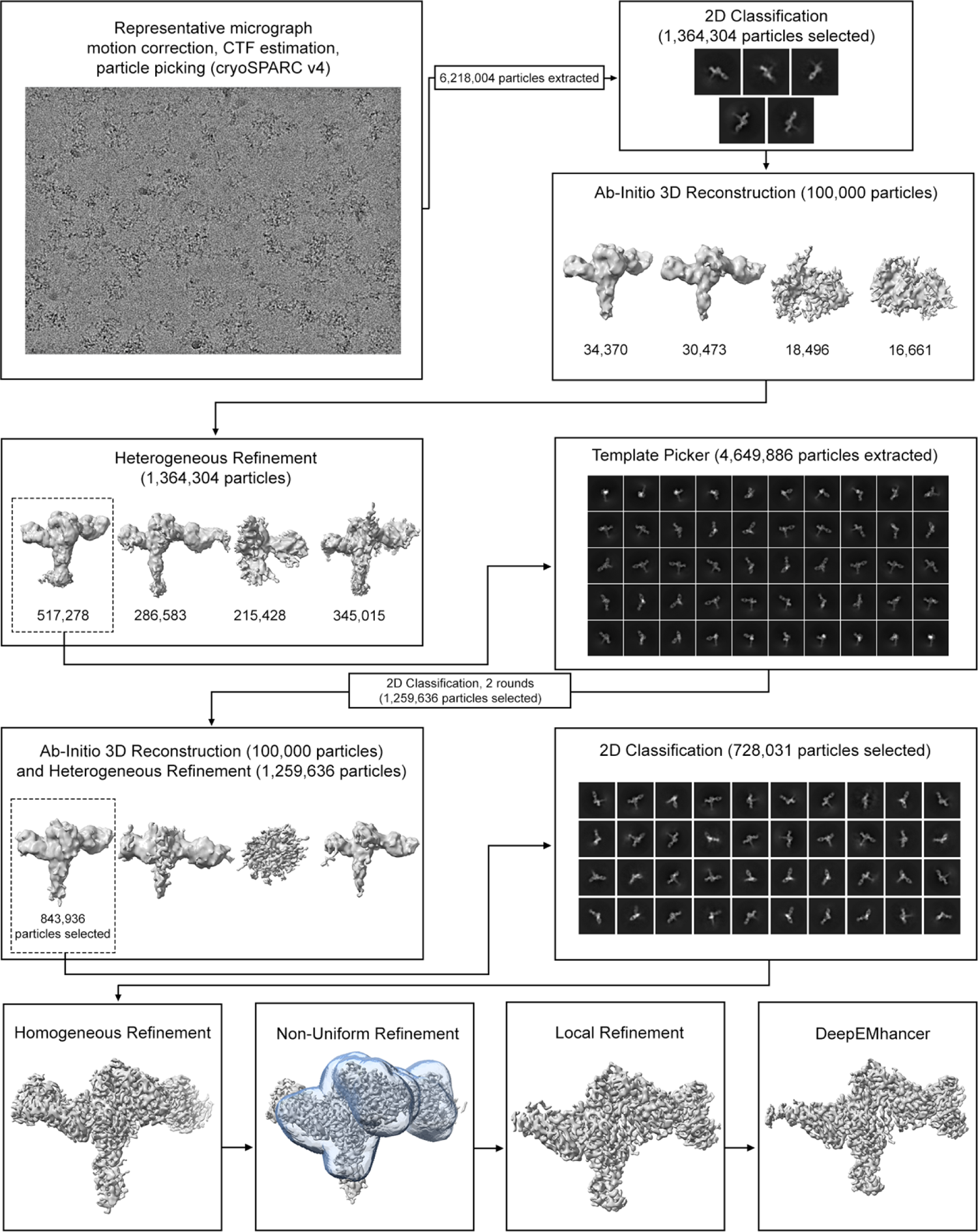
Representative cryo-EM processing workflow of GP38-Gn^H-DS^-Gc. Flowchart outlining cryo-EM processing of GP38-Gn^H-DS^-Gc bound to ADI-46152 and ADI-36125 Fabs.

**Supplemental Figure 6.**
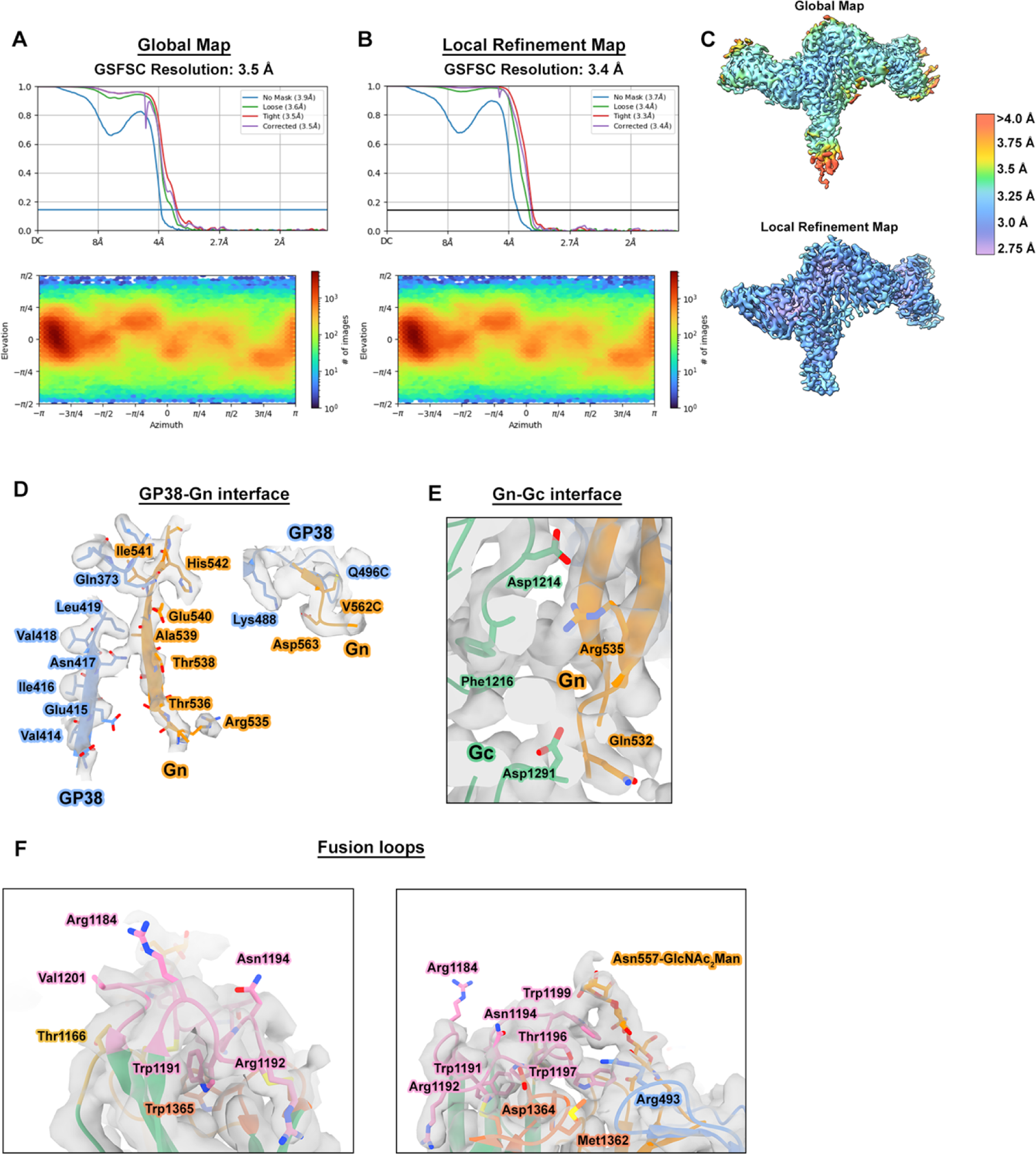
Validation and fit in map for GP38-Gn^H-DS^-Gc. (A) The Gold-standard Fourier shell correlation (GSFSC) curve and angular distribution plot for the global refinement of GP38-Gn^H-DS^-Gc + ADI-36125 + ADI-46152, generated in cryoSPARC v4.0.1. (B) GSFSC curve and angular distribution plot for the local refinement. (C) Local resolution shown by color for the global reconstruction (top) and local reconstruction (bottom). (D-F) Detailed views of the binding interface and corresponding map for the GP38-Gn interface (panel D), the Gn-Gc interface (panel E) and the Gc fusion loops (panel F). Side chains depicted as sticks. Oxygen atoms are colored red, nitrogen blue, and sulfur yellow.

**Supplemental Figure 7.**
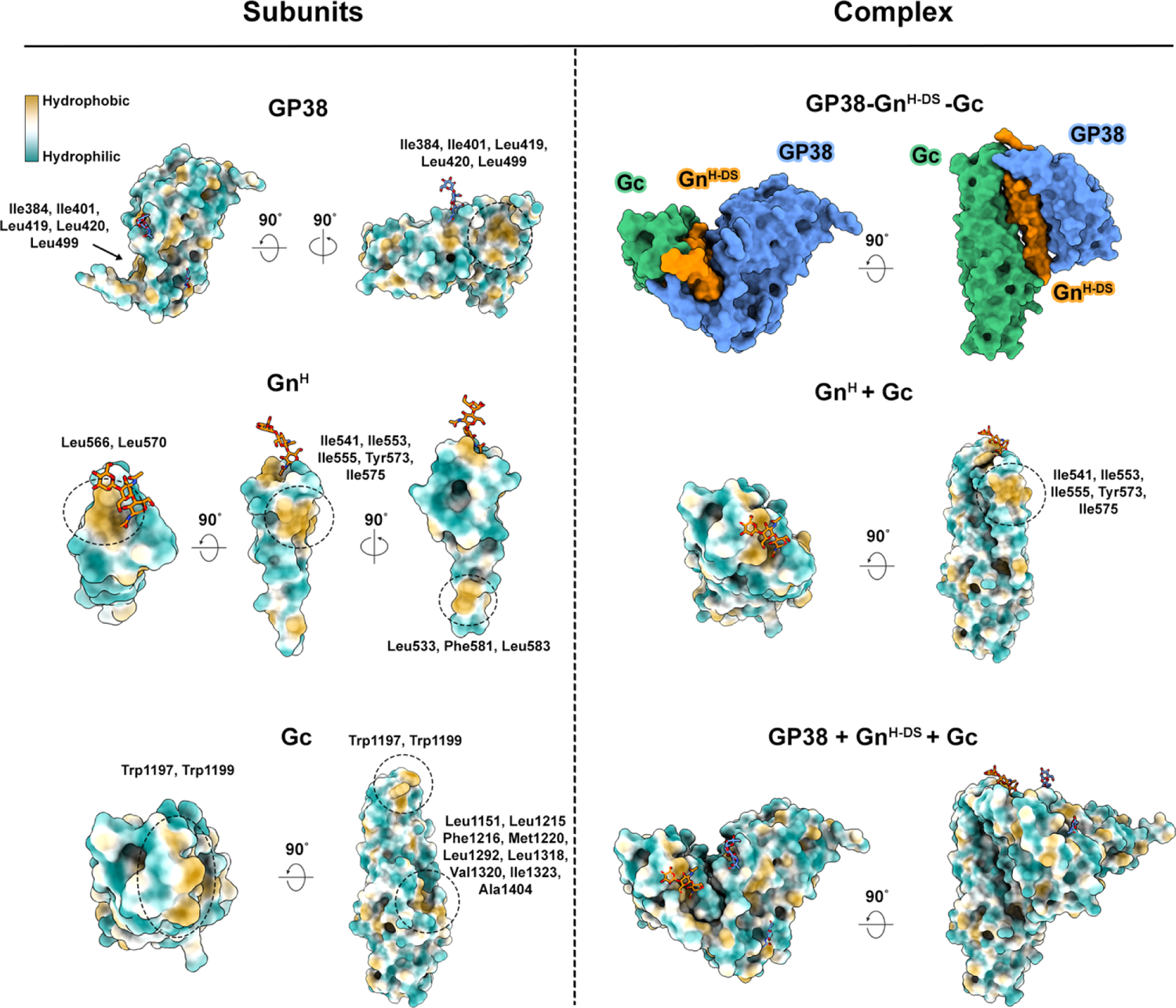
Hydrophobicity analysis of GP38-Gn^H-DS^-Gc. A surface view of components of GP38-Gn^H-DS^-Gc, displaying hydrophobicity. The left half shows the individual proteins. The right panel shows protein complexes. Hydrophobic residues are colored in yellow, and hydrophilic residues are colored in teal.

**Supplemental Figure 8.**
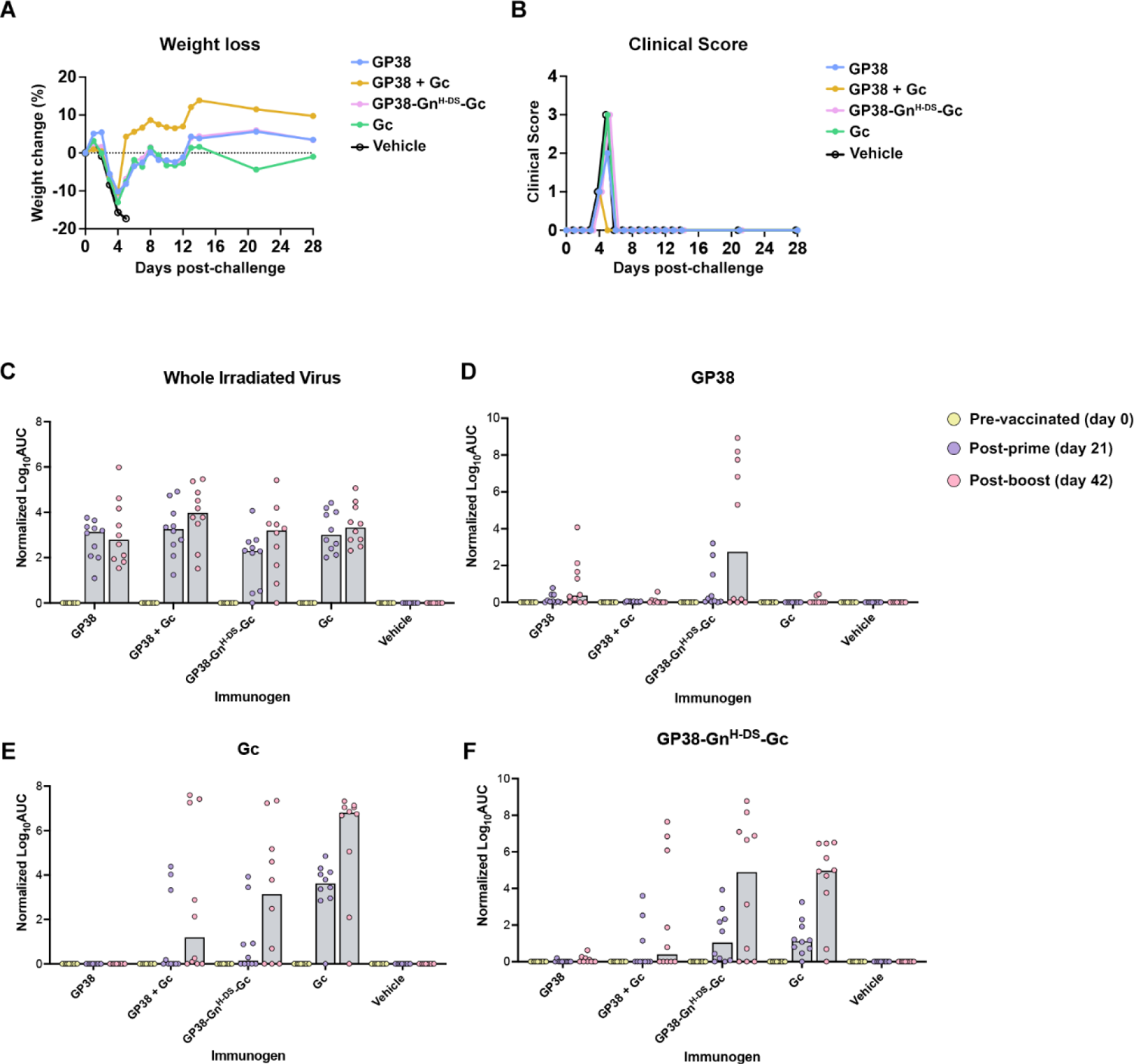
Weight loss, clinical score, and sera binding titers of mice immunized with CCHFV immunogens, related to Figure 7. Values shown are the normalized logarithm of the area under the curve (AUC) for each group. Bar reflects the median value of the individual values, and individual values are shown by points overlayed on the bar.

**Summplemental Table 1.**
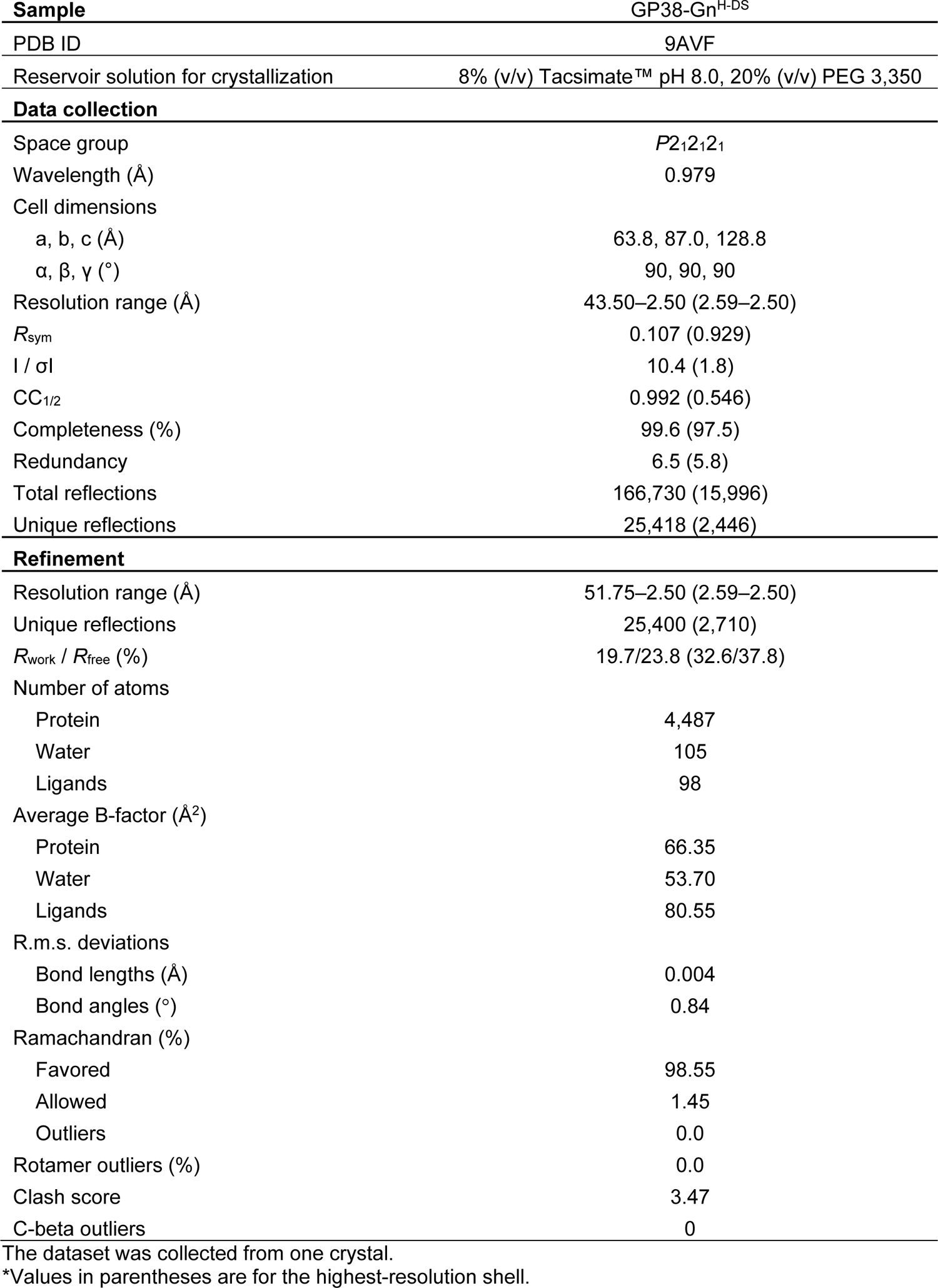
Crystallographic data collection and refinement statistics, related to **Figure 4**.

**Supplemental Table 2.** Cryo-EM data collection and refinement statistics, related to **Figure 6**.

| Sample | GP38-Gn<sup>H-DS</sup>-Gc + ADI-46152 + ADI-36125 |
| --- | --- |
| PDB ID | 9B8J |
| **EM data collection** |  |
| Microscope | FEI Titan Krios |
| Voltage (kV) | 300 |
| Detector | Gatan K3 |
| Energy filter | Gatan Biocontinuum |
| Slit width (eV) | 20 |
| Magnification (nominal) | 105,000 |
| Pixel size (Å/pix) | 0.8332 |
| Exposure rate (e<sup>-</sup>/pix/sec) | 13 |
| Frames per exposure | 100 |
| Exposure (e<sup>-</sup>/Å<sup>2</sup>) | 69 |
| Defocus range (µm) | 1.5-2.5 |
| Tilt angle (°) | 0 (8,061 mics), +30 (5,147 mics) |
| Micrographs collected | 13,208 |
| Micrographs used | 10,739 |
| Particles extracted (total) | 4,649,886 |
| Automation software | SerialEM |
| **3D reconstruction statistics** |  |
| Particles | 728,031 |
| Symmetry | C1 |
| Map sharpening B-factor | -100.4 |
| Unmasked resolution at 0.5 FSC (Å) | 4.4 |
| Masked resolution at 0.5 FSC (Å) | 3.9 |
| Unmasked resolution at 0.143 FSC (Å) | 4.0 |
| Masked resolution at 0.143 FSC (Å) | 3.4 |
| **Model refinement and validation statistics** |  |
| Refinement package | Phenix 1.21 |
| Composition |  |
| Amino acids | 961 |
| RMSD bonds (Å) | 0.004 |
| RMSD angles (°) | 0.91 |
| Average B-factors |  |
| Amino acids | 70.3 |
| Ramachandran |  |
| Favored (%) | 95.94 |
| Allowed (%) | 4.06 |
| Outliers (%) | 0 |
| Rotamer outliers (%) | 0.84 |
| Clash score | 4.79 |
| C-beta outliers (%) | 0 |
| CaBLAM outliers (%) | 2.52 |
| CC (mask) | 0.76 |

